# All roads lead to Rome: Emergence of Mechanistic Divergence Shapes Activation of Class B1 G-Protein Coupled Receptors

**DOI:** 10.64898/2026.01.05.697796

**Authors:** Prateek Bansal, Diwakar Shukla

## Abstract

Class B1 GPCRs are crucial to maintaining homeostasis along a multitude of vital biochemical pathways. Understanding the activation mechanism of these proteins at both a family and clade-specific level is particularly relevant for designing multi-target agonists, as exemplified by recently designed dual-agonists for GLP-1R and GIPR, for treating obesity. Here, we use 6 milliseconds of unbiased all-atom MD simulations of GCGR, GLP1R, PAC1R, SCTR, PTH1R and CALCR from the four different clades of Class B1 GPCRs to establish the universal mechanism of their activation. We show that the activation of Class B1 GPCRs involves a clade-independent intermediate state characterized by the outward movement of helix 6. We use a combination of Markov state models and transition path theory to show that the activation of these proteins occurs at a millisecond timescale. We identify characteristic molecular locks that are conserved at a clade-level, showcasing the uniqueness among the activation mechanisms of these proteins. We show that these proteins show similar inactive and active states, but show unique activation mechanisms at a residue level. These sites can be targeted directly or allosterically to design therapeutics targeting a specific clade of proteins. Thus, this study provides an integrated atomistic view of the activation for Class B1 GPCRs from a mechanistic, thermodynamic and kinetic perspective.

## 1 Introduction

G-Protein Coupled Receptors (GPCRs) form the largest family of cell-surface receptors expressed in the human genome.^1^ These proteins are ubiquitous in human tissue, are involved in maintaining homeostasis across a plethora of biochemical pathways, and are hence critical drug targets.^2^ In response to a drug molecule or endogenous agonists, GPCRs undergo a conformational transition to a stabilized ‘active’ state. This allows the GPCR to bind to G-proteins on the intracellular end, promoting the exchange of GTP for GDP from the G*α* subunit, triggering downstream effects.

On the basis of phylogenetic analysis, these proteins are divided into major classes - A, B(1/2), C, D, F and T. Recent studies show that around 26% of all FDA-approved drugs target GPCRs, with 93% targeting Class A GPCRs^2^ - highlighting the untapped potential of targeting other classes of GPCRs. Class B1 is a sub-family of Class B G-Protein Coupled Receptors consisting of 15 members, which bind peptide ligands.^3^ These are proteins that regulate several crucial biochemical pathways, for example, maintaining blood sugar levels^4^ through the glucagon receptor (GCGR) and the glucagon-like peptide 1 receptor (GLP-1R). Secretin Receptor (SCTR) regulates water homeostasis,^5^ while calcium reabsorption is mediated by via Calcitonin Receptor (CALCR) and Parathyroid Hormone - 1 receptor(PTH1R).^6^ Additionally, stress response is mediated via Pituitary Adenylate Cyclase Activating Polypeptide - 1 Receptor (PAC1R).^7^ These receptors have an extracellular domain of ∼ 120 residues, which is important for sensing peptide agonists in addition to the prototypical heptahelical transmembrane domain of ∼ 260 residues.

A considerable amount of literature exists on the activation of Class B1 receptors, which has been studied in detail, through structural studies that resolve high-resolution structures of these proteins.^8–11^ Although these structural studies give us an idea about the inactive and active states of these receptors, we do not have a clear picture of the intermediate steps that take place during activation, which might involve states critical for targeted drug design.^12–15^ Additionally, we posit that, since these proteins share high structural similarities in their active and inactive states (Fig. 1a), there might be unresolved universal links among the activation mechanisms of these proteins.

**Figure 1:**
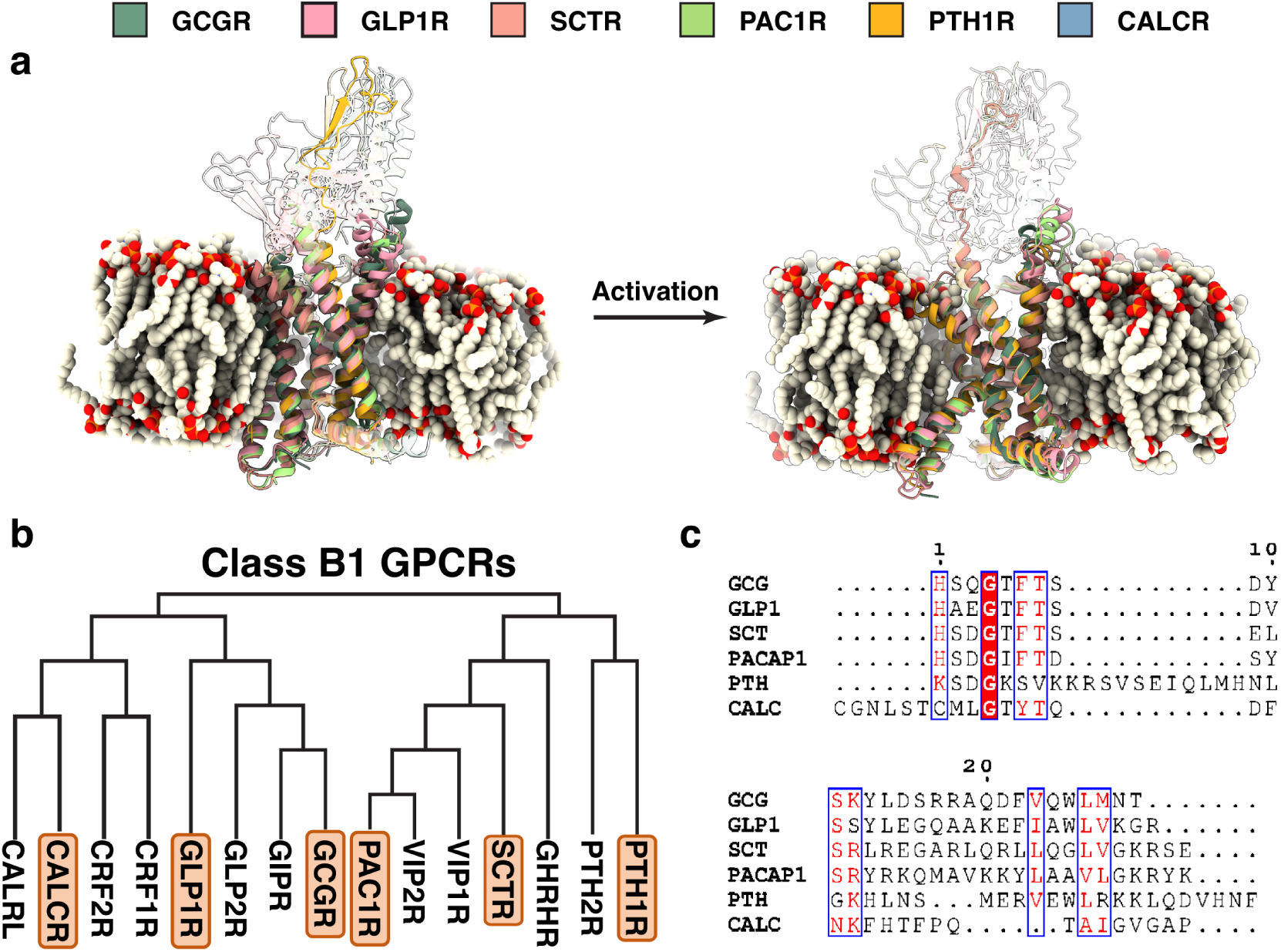
Activation in Class B1 GPCRs. (a) Activation processes in Class B1 GPCRs, shown by the intracellular kinking of TM6. Six proteins used in this study GCGR (dark green), GLP1R (salmon), SCTR (orange), PAC1R (light green), PTH1R (yellow) and CALCR (blue). Each protein was embedded in a membrane and simulated to uncover the entire activation pathway. (b) Phylogenetic tree representing the Class B1 of GPCRs. Proteins chosen for this study are highlighted in orange. (c) Sequence alignment among the different Class B1 endogenous peptide agonists.

Despite their therapeutic relevance, Class B1 GPCRs have until recently remained relatively understudied as compared to the class A GPCRs due to their complex peptide-mediated activation that involves binding of peptides to a flexible extracellular domain. The substantial conformational heterogeneity of class B GPCRs needed more sophisticated crystallographic and Cryo-EM techniques as compared to Class A GPCRs. ^16^ Mechanistically, both Class A and B1 GPCRs show an outward movement of TM6 upon activation, but this analogy does not extend to residue-level movements. The conserved NPxxY/DRY motifs have been implicated in stabilizing the inactive state for Class A receptors, but are absent in Class B1 GPCRs.^17^ Instead, the conserved PxxG motif is implicated in promoting the formation of a sharp kink in TM6 (typically ∼90–100*^◦^*).^18^ The existence of the ECD-TMD binding site, as well as Class B1 specific microswitches can explain why Class B1 activation has been less tractable and provides a basis for studying the activation of these unique proteins.

The activation mechanisms of Class B1 GPCRs have been outlined by several studies, using Cryo-EM structures, mutagenesis and activity assays. The first such structure of a Class B1 GPCR (GLP-1R) in the active state^9^ showed a ∼ 18 Å outward movement of the intracellular end of TM6. This large movement allows for G-protein engagement at the intracellular end. Cryo-EM and DEER spectroscopy show that GCGR undergoes a sharp kink and a helical break in TM6 in the presence of G-protein, and that the barrier for activation is much higher in the absence of an agonist.^19^ This is in contrast to Class A GPCRs, where TM6 undergoes bending, but the helicity is preserved.^20^ In the case of CALCR, using phase-plate cryo-EM, a Gs bound structure revealed a 60*^◦^* kink in helix 6 to accommodate G*α*’s C-terminus. Similarly, the calcitonin gene-related peptide (CGRP) receptor requires TM6 kinking around P^6.47*b*^ (Superscripts indicate the Wootten numbering system^21^ introduced in for Class B1 GPCRs) for activation and mutation of this proline abolishes the kink and impairs signaling.^22^ In the PTH1R-Gs structure, this kink angle of TM6 with the core of the protein is closer to 90*^◦^*. This kink is absent in the antagonist bound state.^23^ SCTR, in addition to the characteristic outward kinking, also shows unique interactions with the peptide - showing that Class B1 GPCRs show varied binding modes with peptides.^24^ For the PAC1 receptor (PAC1R), cryo-EM structures indicate that peptide agonist binding destabilizes TM6 around the PxxG motif, making it bent upon G protein engagement. Mutating T^6.42*b*^ of the HETx or P^6.47*b*^ PxxG motif significantly affects activity. ^25,26^ Additionally, mutation of G^6.50*b*^A stabilizes GLP1R,^10^ with G^6.50*b*^C being used for crystallizing GLP1R.^27^ Hence, there is strong evidence for TM6 bending being a proxy for receptor activation in Class B1 GPCRs. The question here, is whether the pathways taken by these proteins to undergo the transition between the inactive and active structures are similar or distinct.

Receptors in this family are important drug targets for diabetes, with recent advances including semaglutide ^28^ (Ozempic) and tirzepatide^29^ (Mounjaro). Multi-receptor agonists, such as dual incretin agonists peptide15, cotadutide, and SAR425899^30^ which target both GCGR and GLP1R, as well as the triple agonist retatrutide ^31^ (GCGR, GLP-1R, GIPR) show conserved peptide-protein contacts via common motifs. The development of these multi-protein agonists further supports a clade-specific basis of activation among these proteins, an aspect that has yet to be explored using dynamic simulation approaches. Although multiple computational studies have employed biased sampling methods, ^32,33^ such as meta-dynamics^33,34^ and Gaussian-accelerated molecular dynamics,^32^ these approaches have limitations. These studies, however, do not provide insights into the activation mechanism in the absence of the peptide - a measure of the basal activity of these proteins. Here, we use unbiased MD simulations to delineate the entire activation process in atomistic detail. Moreover, a recent study used a large-scale genome-wide MD simulation dataset highlighting the opening of allosteric sites and lateral gateways on these receptors at microsecond timescales underscoring the need for large-scale MD simulation studies to understand the dynamics of these integral proteins.^35^

Here, we present the insights from more than 6 ms (Table S1) of performed unbiased MD Apo + Holo simulations, used to study the activation mechanisms of Class B1 GPCRs as a whole. Additional simulations were performed in the presence of endogenous agonists, to investigate the allosteric pathways for ligand-based activation. Markov State Models were used to describe the activation mechanisms of these proteins, to resolve the thermodynamics and kinetics of the activation process. Recent studies have successfully outlined the activation mechanisms of GPCRs using Markov state models, ^13,32,36–42^ thereby facilitating selective drug discovery.^13,43^ We show that the activation mechanisms of these proteins show a stage-wise conserved outward movement of TM6, followed by kinking at the intracellular end of TM6 - a hallmark of GPCR activation. We show that the activation mechanisms are conserved on a clade-specific level, rather than a family-wide universal activation mechanism. Additionally, we identify motifs that are conserved across the entire family, unique to each clade, as well unique to each protein - which play a role in activation. This gives us a basis to understand the activation mechanism from three levels: across the entire family, within specific clades and at the level of individual proteins.

## 2 Results and discussion

### Class B1 GPCR activation is marked by a conserved outward movement and intracellular kinking of TM6

To study the activation processes of these proteins in detail, we started with the active and inactive states of each protein embedded in a lipid bilayer (Fig 1a). Six of the fifteen Class B1 GPCRs - Glucagon receptor (GCGR), Glucagon-like peptide 1 receptor (GLP1R), Parathyroid Hormone 1 Receptor (PTH1R), Calcitonin Receptor (CALCR), Secretin Receptor (SCTR) and Pituitary Adenylyl Cyclase Activating Polypeptide 1 Receptor (PACAP-1R) - belonging to all the major subclades of the family (Fig. 1b) were chosen as the candidates. These proteins had resolved structures including the ECD. Simulations were performed starting from both the active and inactive states, until transitions from the inactive to active and vice-versa were observed. To accelerate sampling, an adaptive sampling strategy was used where the simulations were carried out in rounds. ^44,45^ At each round, the trajectories were featurized and clustered, and frames belonging to the least populated clusters were used to seed the next round of simulations. To correct for the sampling bias introduced by sampling the rare states more frequently, Markov State Models were constructed to get an unbiased representation of the thermodynamic and kinetics of the activation process. MSMs have been routinely used to analyze and reweigh the conformational ensembles of GPCRs,^38,40^ and here we employ them to analyze the dynamics of an entire family of GPCRs. The microstates for MSM construction were defined by clustering on a reduced dimensional space using time-lagged indepedent component analysis (tICA) (Fig. S1), enabling the capture of the slowest kinetic processes.

Overall, activation and deactivation was observed for all proteins, using the metrics previously used to describe canonical Class B1 GPCR activation - the *ϕ* backbone dihedral angle at G^6.50*b*^, which captures the kinking of TM6. This was plotted against TM3-TM6 distance measured at the intracellular end, which captures the outward movement of TM6, to accommodate the G-protein. These order parameters were introduced by Mattedi et al., 2020. Upon projection of the dataset of each protein along these reaction co-ordinates and computing the free energy associated with each frame, we observe the entire activation pathway for each protein (Fig. 2). Despite the conserved topology in the inactive and active states, each receptor showed unique activation pathways. In each receptor, the active and inactive states were differentiated by at least one intermediate state. In the case of GCGR, the intermediate state observed in GCGR is similar to the one discussed previously, ^34^ where TM6 is displaced outward but remains un-kinked. This provides a validation for our work, and we are able to identify structures from which simulations were not started. For each receptor, the activation pathway consists of the outward movement of TM6, followed by kinking at G^6.50^ to reach the final active state. The outward movement of TM6 consists of a large barrier of ∼ 2.5-3 kcal/mol for all proteins. However, the highest barrier is the kinking of the helix, which is often ∼ 4-4.5 kcal/mol for all proteins. This is in line with the highest barriers observed along tIC1 (Fig. S1).

**Figure 2:**
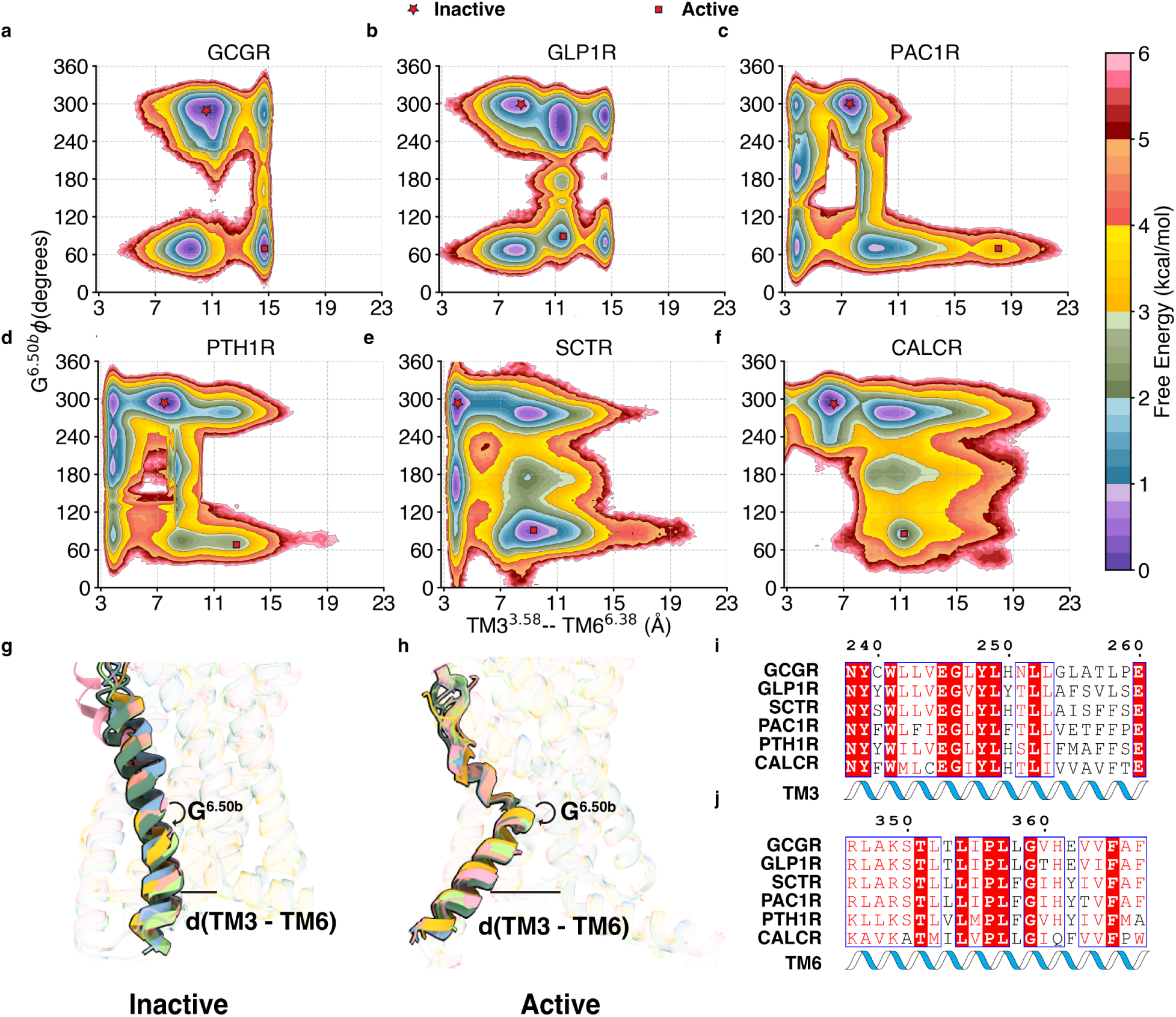
Overall Activation of Class B1 GPCRs. (a-f) Free energy landscapes showing the stage-wise outward movement of TM6 in the six proteins considered in this study. The backbone dihedral *ϕ* at *G*^6.50*b*^ was plotted against the distance measured between residues L^3.57*b*^ and T^6.42*b*^ for (a) GCGR (b) GLP1R (c) PAC1R (d) PTH1R (e) SCTR and (f) CALCR. (g-h) Visual schematic showing the difference between (g) inactive and (h) active states for each of the six GPCRs. (i-j) Multiple sequence alignments showing the conservation of residues L^3.57*b*^ and T^6.42*b*^.

Interestingly, even though these proteins show similar inactive and active states (Fig 1b), we observe that the intermediate states are present at different points in the tIC landscape (Fig. S1). To further compare the unique intermediate states observed, we computed the Kullback-Liebler Divergence for each residue pair. Upon projecting the KL divergence values on the backbones of these proteins, we observe that GCGR and GLP1R show similar intermediate states, SCTR and PAC1R show similar intermediate states, while PTH1R and CALCR show distinct unique intermediate states (Fig. S2, S3). These differences are concomitant with the phylogenetic arrangement of these proteins within the Class B1 family (Fig. 1c). Hence, there exists a strong evidence for a clade-wise unique activation mechanism for these proteins. While previous studies have focused primarily on static end-states of Class B1 GPCRs, our unbiased simulations resolve the entire activation trajectory, including high-energy intermediates that are difficult to capture via experiment. Crucially, we observe that these intermediate states differ across subclades, consistent with the phylogenetic clustering of Class B1 GPCRs. This provides evidence for a clade-specific activation mechanism, advancing our understanding beyond the conserved endpoint structures reported in previous cryo-EM studies.^23,24,26^

### Class B1 G-Protein Coupled Receptors show similar active states but distinct activation mechanisms

To further drive into the details of activation from a clade-wise perspective, we posited that the differences in activation mechanisms are due to the unique dynamic contacts that are broken and formed during the process of activation. We hence projected our data into structural metrics that were considered important for a specific clade - residues that are only conserved in a certain clade and not conserved in other proteins within the family. Class B1 GPCRs can be divided into two superclades - one containing GCGR, GLP1R and CALCR, while the other containing SCTR, PAC1R and PTH1R (Fig. 1b). For the clade containing GCGR and GLP1R, we observed that the intermediate state shows a major movement in the extracellular end of TM7 between the inactive and intermediate states (Fig S2). Hence, we sought to find locks that were broken during activation that were unique to GCGR and GLP1R. First, a multiple-sequence alignment for all receptors under consideration was generated, and residue pairs where at least one residue that was conserved on a clade-specific level were identified. These residue pairs were then filtered for physical proximity within the receptor and for the potential for form polar interactions, which could form or break during the activation process. To confirm if these locks were broken concomitant with other indicators of activation, we produced a realization of the MSM using Kinetic Monte Carlo simulations - yielding GPCR activation trajectories (Fig. 3 a). These trajectories were then used as a proxy for studying the overall activation process (Fig. 3a-e).

**Figure 3:**
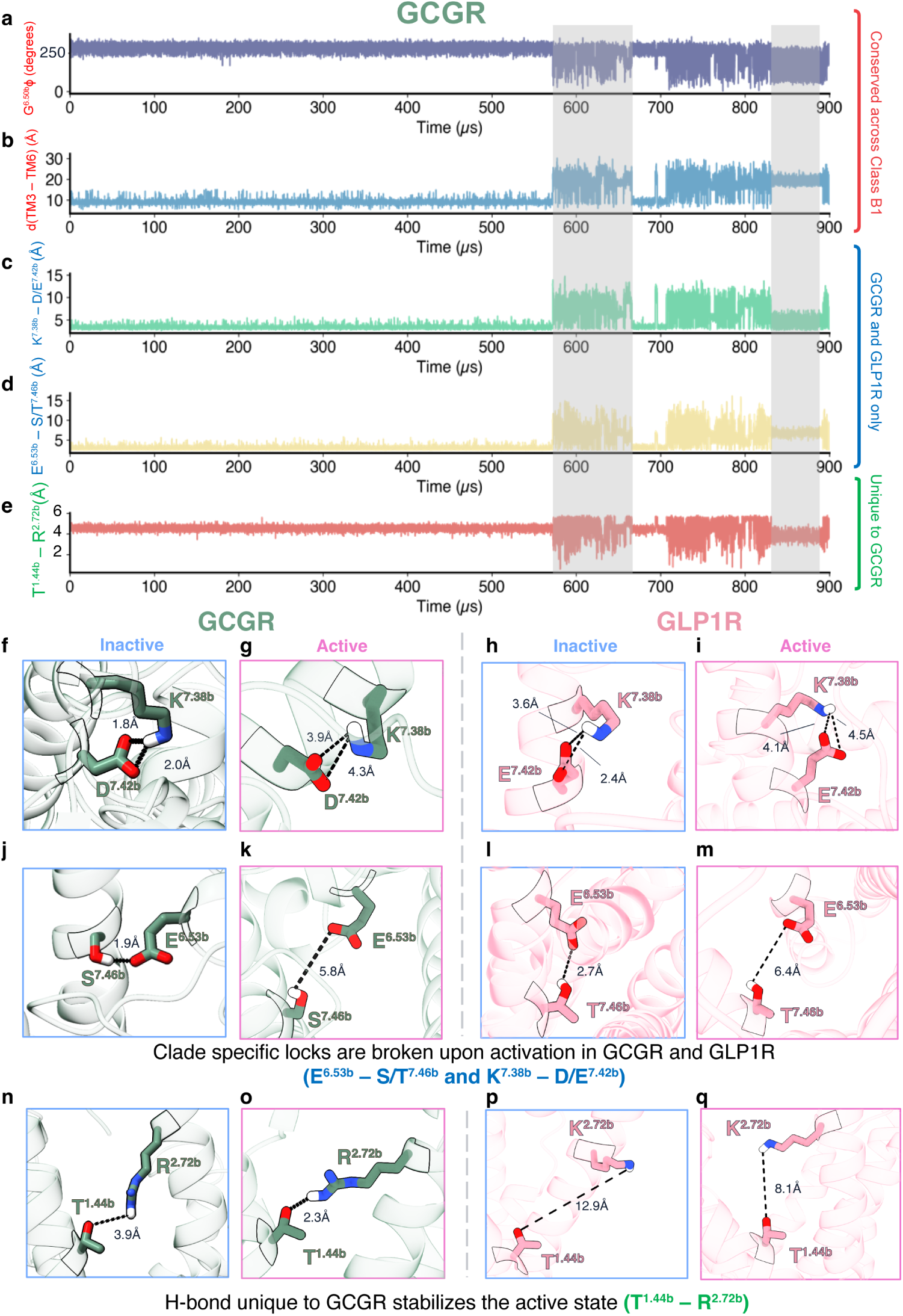
Clade-wise activation mechanism in the GCGR-GLP1R clade. (a-e) KMC simulations performed for GCGR. Active confirmations are highlighted in light grey. The plots show (a) backbone dihedral angle *ϕ* at G^6.50*b*^, (b) distance between TM3 and TM6 measured at L^3.57*b*^ and T^6.42*b*^, (c) distance between K^7.38*b*^ and D/E^7.42*b*^ (d) distance between E^6.53*b*^ and S/T^7.46*b*^ and (e) distance between T^1.44*b*^ and R^2.72*b*^ for GCGR. (f-i) salt bridges between K^7.38*b*^ and D/E^7.42*b*^ break as the proteins transition from inactive (f,h) and active (g,i), for GCGR (f,g) and GLP1R (h,i). (j-m) hydrogen bonds between E^6.53*b*^ and S/T^7.46*b*^ break as the proteins transition from inactive (j,l) and active (k,m), for GCGR (j,k) and GLP1R (l,m). (n-q) (f-i) hydrogen bond between T^1.44*b*^ and R^2.72*b*^ forms for GCGR (n,o), but not for GLP1R (p,q).

On projecting the activation metrics as discussed in the previous section - the intracellular distance between TM3 and TM6 as well as the dihedral angle *ϕ* at *G*^6.50*b*^ (Fig. 3a,b), we see that activation occurs when the distance between TM3-TM6 is high (between 10 and 15 Å) and and the dihedral angle is low (between 0*^◦^* and 60*^◦^*). This is highlighted as the light gray section in the plot. Interestingly, our data shows that the breaking of a salt bridge (K^7.38*b*^-D^7.42*b*^ in GCGR and K^7.38*b*^-E^7.42*b*^ in GLP1R) is correlated with protein activation for these two proteins (Fig. 3 c,f-i). Additionally, we also observe a hydrogen bond (E^6.53*b*^-S^7.46*b*^ in GCGR, E^6.53*b*^-T^7.46*b*^ in GLP1R) shows similar behavior on protein activation (Fig. 3 d,j-m). These motifs are not observed in any other protein as they are not conserved in the other members of Class B1 GPCRs. There is another molecular lock specific to GCGR only - between T^1.44*b*^ and R^2.72*b*^ that is broken on activation. This lock is not present in GLP1R, as R^2.72*b*^ is replaced by K^2.72*b*^, and the ammonium group of K^2.72*b*^ cannot insert into the core of TMD, as afforded by the guanidium group of R^2.72*b*^. These locks are specific to the clade, and hence targeting these locks directly or allosterically would be an effective strategy for designing drugs that target multiple members of this family.

For the second clade containing SCTR, PAC1R and PTH1R, we interestingly observe clade-specific locks that are formed on activation, possibly to stabilize the active state (Fig. S4). For this case, GPCR activation trajectories were constructed as well (S4a-d). A hydrogen bond between the sidechain of S^6.59*b*^ and the backbone oxygen of F^6.56*b*^ is formed (S4c,e-j) for SCTR (S4c,e,f), PAC1R (S4g,h) and PTH1R (S4i,j). For the clade containing SCTR and PAC1R but excluding PTH1R, we also observe a hydrogen bond (R^5.40*b*^-E*^ECL^*^3^ in SCTR, K^5.40*b*^-E*^ECL^*^3^ in PAC1R) shows similar behavior on protein activation (Fig. S4 d,k-n). The formation of these hydrogen bonds indicates a conserved mechanism for stabilizing the active state in this superclade.

Taken together, our findings illustrate that Class B1 GPCR activation involves both universal and clade-specific molecular rearrangements. The identification of unique dynamic interactions, such as clade-specific hydrogen bonds and salt bridges, provides essential mechanistic insights into the within this receptor family. These interactions are unique to the specific clade, and enables future targeted therapeutic design.

### Activation of Class B1 GPCRs occurs over a millisecond timescale

In the above section, we have focused on the thermodynamic perspective of activation of Class B1 GPCRs. Kinetic timescales of activation are a crucial indicator of the dynamic accessibility of the active state and potential bottlenecks in signal transduction. To emphasize on this, we computed the average timescales of activation for Class B1 GPCRs using the Markov State Models and Transition Path Theory. To compute the timescales, the source (inactive) and destination (active) microstates were chosen from the state decomposition. Then, reactive trajectories were computed that started at the inactive microstate and ended in the active microstate.

For Class B1 GPCRs, we observe that the activation process can be broken to two steps - from the inactive to the intermediate state, and then from the intermediate to the active state. The first transition from the inactive to intermediate state (characterized by the straight outward movement of TM6) occurs on an average timescale of 325 - 450 *µ*s (Fig. 4a,b). This is followed by the intermediate to the active state (characterized by the breaking and kinking of TM6), occurring on a longer timescales - from 550 - 900 *µ*s (Fig. 4b,c). If the direct transition from the inactive to active state is considered, the activation process for each Class B1 GPCR considered in this study occurs on a 780 - 1400 *µ*s. This means that the flux of transitions from inactive state to the active state must past through the transition state - since the timescales of overall activation are similar to the timescales of the two intermediate steps added.

**Figure 4:**
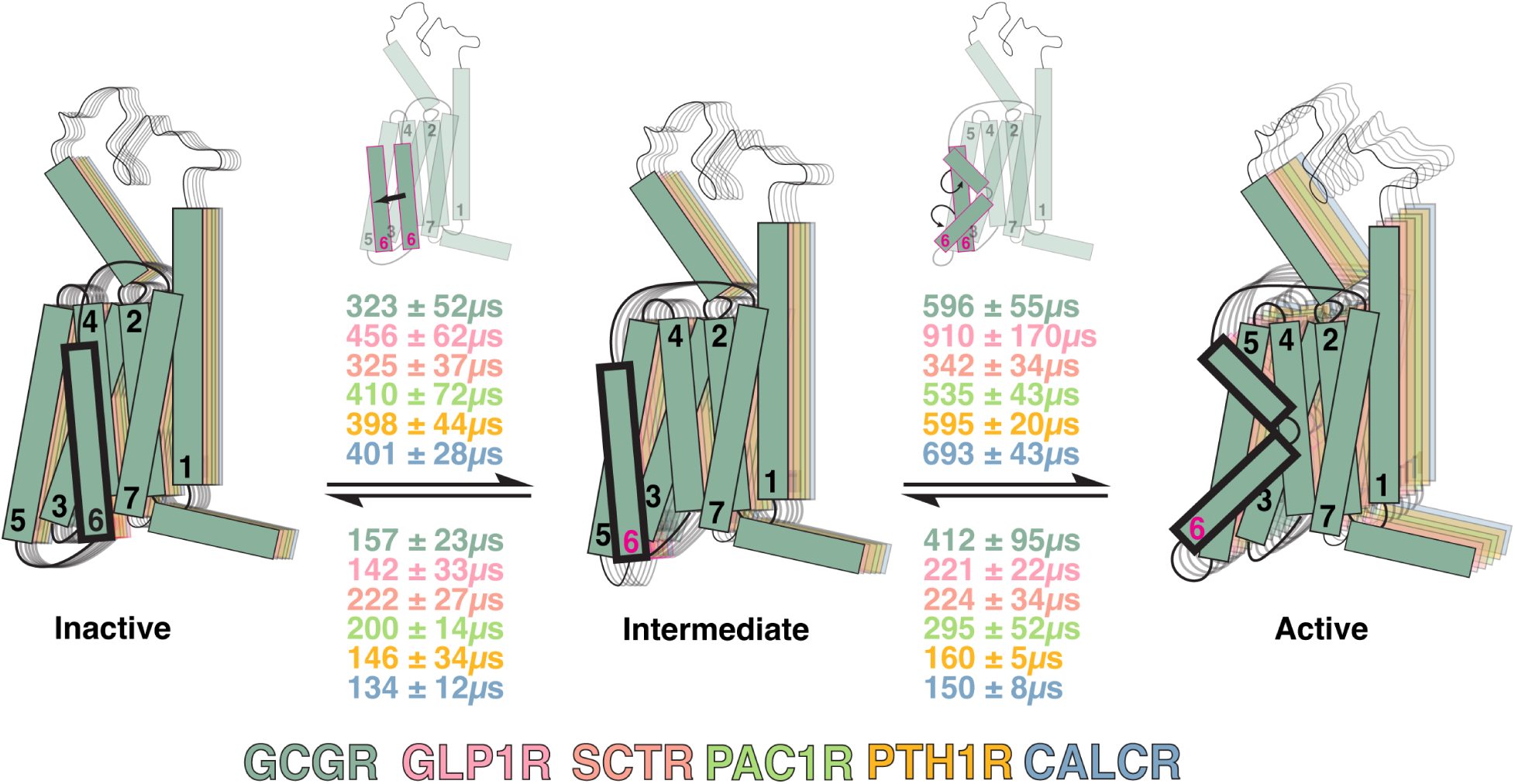
Timescales of activation in Class B1 GPCRs. (a) Combined inactive state (b) Combined intermediate state (c) Combined Active state for each GPCR considered in this study. Transition between (a) and (b) is characterized by the straight outward movement of TM6 (bold) whereas the transition between (b) and (c) is characterized by kinking and breaking of TM6 (bold). Timescales of activation are presented for the forward and reverse processes. Each transition timescale is colored coded with the proteins.

**Figure 5:**
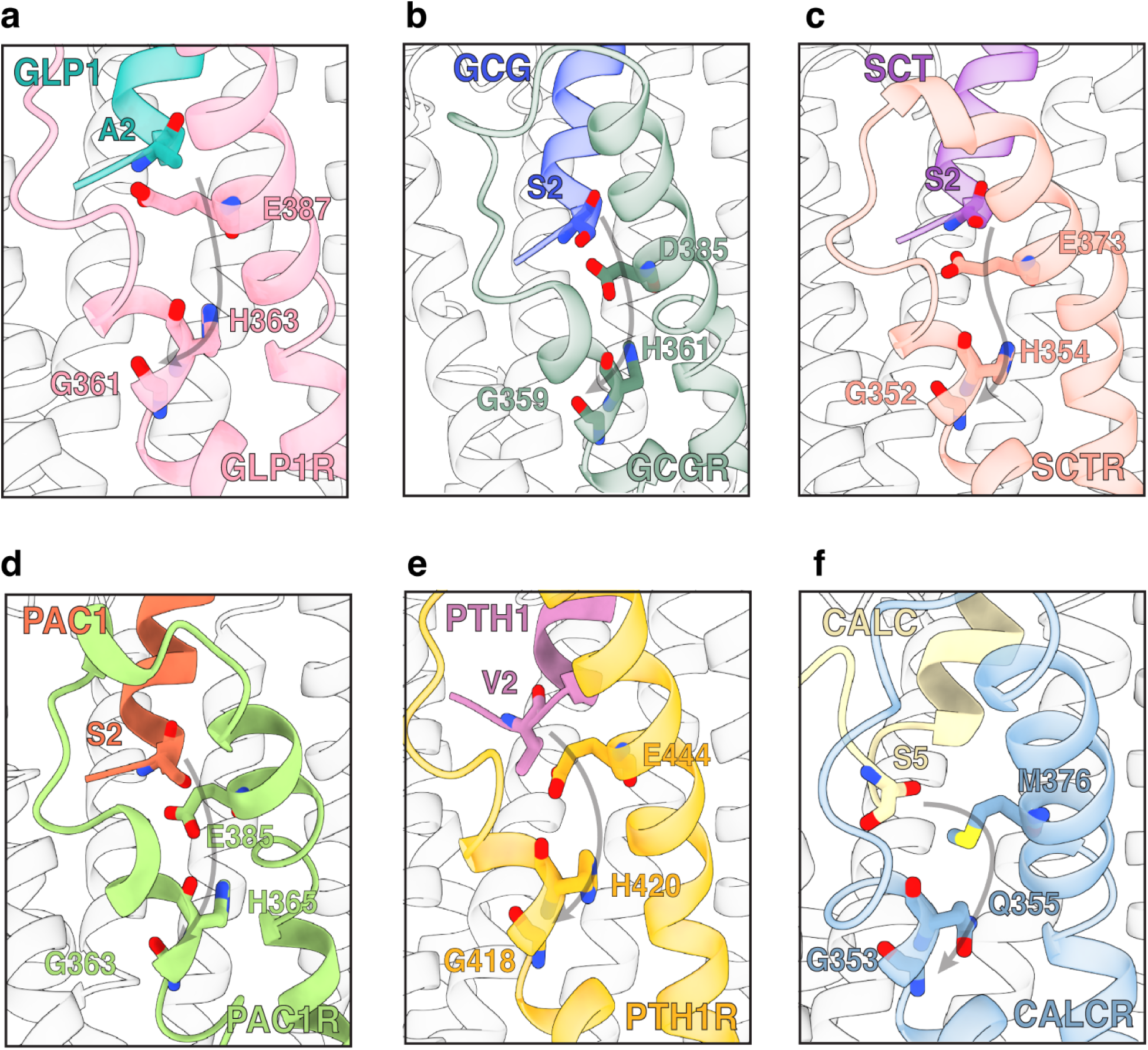
Conserved Allosteric pathways in Class B1 receptors. Pathway begins at the 2nd position of the peptide (S/A2), followed by conserved residues D/E^7.42*b*^ and H^6.47*b*^, and ending at G^6.50*b*^. Shown are the allosteric pathways for (a) GLP1R-GLP1, (b) GCGR-GCG, (c) SCTR-SCT, (d) PAC1R-PACAP, (e) PTH1R-PTH and (f) CALCR-CALC receptor-peptide complexes.

The timescales associated with the reverse process are much faster - showing that the deactivation of these proteins has a smaller kinetic barrier than the activation process. This makes the overall activation process a rare process, necessitating the use of long timescale MD simulations to characterize them. These findings highlight that the formation of the kinked TM6 conformation is the main kinetic bottleneck in the activation of Class B1 GPCRs, consistent with experimental observations that G protein engagement is essential for full activation. Furthermore, the faster deactivation kinetics imply that these receptors are intrinsically tuned for rapid signal termination, potentially to ensure precise temporal control in physiological signaling. Together, our results provide a kinetic framework for understanding the timing and efficiency of signal transduction in Class B1 GPCRs.

### Conserved allosteric pathway in Class B receptors mediates the signal from the peptide to the PxxG motif

Class B1 GPCRs in the recent years have seen a significant rise in the number of peptide bound structures which has allowed for a detailed look into the peptide-based activation mechanism of these proteins. These receptors share a conserved peptide binding mode - peptide agonists occupy the binding site at the ECD-TMD interface. This is enabled by the C-terminus of the peptide inserting itself into the core of the TMD and the N-terminus engaging with the ECD.^16^ Studies have shown that agonist binding reduces the barrier for the outward movement of TM6 - leading to G-protein binding.^16^ However, the interesting question is that the binding site of the peptide lies 20 Å away from the G-protein engagement site - suggesting that there is an allosteric communication between the peptide and the G-protein engagement site.

To identify this universal effect by which Class B1 peptides modulate conformational change in their respective receptors, we calculated allosteric pathways that link the peptide to the G-protein engagement site. Each allosteric pathway started at the second residue - a residue that is semi conserved across all peptides. Since the bending occurs at a conserved glycine in the PxxG motif - we chose G^6.50*b*^ as the end point of each pathway. An allosteric pathway connects conformationally coupled residues that show correlated fluctuations. These pathways form the basis of communication within proteins, an essential feature of GPCRs. Studies on class A GPCRs show a crucial role for these pathways in signaling, with disruption of these pathways considered a primary reason for reduced function in GPCRs.^46^

The pathways were computed using the peptide-bound simulations of each of the six proteins. Mutual Information was used to compute the flux of information through the protein as a graph - with the nodes being the residues and the edges being the strength of allosteric coupling between a pair of residues. For each pathway, the second residue of the pathway was the conserved residue D/E^7.42^, in the receptor, which has been known to interact at position 2 of the peptide.^47,48^ Interestingly, D/E^7.42^ also forms a clade-specific lock with K^7.38*b*^ in the clade containing GCGR and GLP1R, further underscoring the functional importance of this residue. Computed allosteric pathways further corroborated this interaction as crucial for passing the signal from the peptide to the protein. Mutation of residue D/E^7.42^ (refs. ^49–52)^ has been shown to affect ligand binding and intracellular signaling. The third residue along the pathway, a partially conserved H^6.52^, also involved in the pathway, a residue that has been shown to be mutationally intolerant for surface expression.^21,50^ This shows us that Class B1 GPCRs use a conserved coupled pathway to relay signals from the peptide to the site of G-Protein engagement. Thus, the mechanism through which endogenous peptides effectively modulate Class B1 GPCR activity is established, showing an integral role for TM6 and TM7 in signal transduction.

## 3 Conclusions

In this study, we show that the overall activation of these proteins follows elements of universal change while also maintaining distinct clade specific changes. Using a combination of Markov State Models and long-timescale Molecular Dynamics Simulations, we outline the activation process of six Class B1 GPCRs. We show that the outward movement of the intracellular end of TM6 occurs in a stage-wise process with the unkinked helix moving outward, followed by kinking. We show that the activation process involves transitions between the inactive and active states - which are similar for all proteins. However, the intermediate states are distinct - showing that there is more to activation of these proteins than just the intracellular kinking of TM6.

We show that GCGR and GLP1R show structural changes specific to the clade - a salt bridge and a hydrogen bond that are broken on activation of these proteins. These motifs are not conserved in other proteins of Class B1 GPCRs, and hence provide a platform to target these proteins for drug discovery. Thus, using *>* 6 ms of simulations, we resolve a unifying yet clade-aware blueprint for Class B1 GPCR activation. We show that these receptors show similar inactive and active states, but distinct activation mechanisms.

We also show that the activation of these proteins occurs on the hundreds of microseconds timescale, and that the majority of the flux of the activation process passes through the intermediate state. Using mutual information calculations, we show that the peptides communicate with the proteins using a conserved allosteric pathway, giving insights into ligand-based activation of these proteins.

One of the limitations of the study is that we are not able to comment on the overall ligand-based activation process for these proteins, which is reserved for future work. Insights from our study can help shed light into the previously unknown sites of targeting these proteins, and hence create newer avenues for drug discovery.

## 4 Methods

### 4.1 Molecular Dynamics Simulations

#### 4.1.1 System Setup

##### 4.1.1.1 Apo systems

For studying the activation of various Class B1 GPCRs, MD simulation systems were setup for each protein, in the inactive and active states, without the presence of the endogenous ligand (Apo). For each protein, the inactive and active conformations were taken from the Protein Data Bank (PDB). PDB IDs of each of the initial structure used to setup a simulation system are provided in Table S2. For each system, the bound ligand and stabilizing antibodies (if present), were removed. Mutations performed to the protein for efficient structure determination were reverted back to Wild-Type Mutation details are provided in Table S3. The missing residues in each structure were modelled using MODELLER (Table S4). For systems that didn’t have PDB structures for one of the conformations, the Alphafold2-Multistate prediction for the specific protein was used as the starting point instead. These structures were obtained from GPCRdb.^11,19,24,26,27,53–59^ Details for such systems are provided in Table S2. Thus, in total 12 systems (6 Inactive + 6 Active) for each of the 6 proteins were prepared. Protonations of each of the acidic residues (H/D/E) were determined at physiological conditions using the H++ server.^60^ Details of the protonations added to each system are provided in Table S5. Protonations were kept across the active and the inactive starting point for each system. Termini of each protein were capped using neutral caps - ACE for N-termini, and NME for C-termini. Each protein was embedded in a POPC bilayer using CHARMM-GUI.^61^ Atomic interactions were characterized using the CHARMM36m force field.^62^ Each system was solvated using TIP3P water^63^ and 150 mM NaCl, to mimic physiological conditions. System sizes for each system are reported in Table S6. Non-protein hydrogens were repartitioned to 3.024Da to facilitate stable dynamics at a higher timestep (4fs).^64^

##### 4.1.1.2 Holo systems

For computing the allosteric pathways associated with activation, each GPCR was also simulated in the presence of the endogenous peptide agonist (Holo). System setup for Holo systems is identical to the Apo systems for the receptors. For the holo systems, the proteins were in the active confirmation. The peptides were capped with neutral termini. Protonations of the peptides were computed using H++ server. ^60^ None of the peptides showed any protonated acidic residues at endogenous pH. Total 6 systems (one for each protein) were prepared. System details are mentioned in Table S2.

#### 4.1.2 Pre-Production Simulations

Each of the 18 systems (12 Apo + 6 Holo) were subjected to minimization for 15000 steps, using the steepest descent method. This was followed by minimization for 20000 in the presence of restraints on the hydrogen bonds, using the SHAKE algorithm for non-water molecules and SETTLE algorithm for water molecules.^65^ Systems were then heated to 310 K using the NVT ensemble, over 10 ns each. For the first 3.1 ns, the overall temperature was increased linearly from 0 to 310 K, followed by keeping the temperature constant for 6.9 ns. During this procedure, the protein backbone was harmonically constrained using a force constant of 10 kcal·mol*^−^*^1^·Å*^−^*^2^. Systems were then equilibrated using the NPT ensemble at 1 bar for 10 ns, using the NPT ensemble, again in the presence of the harmonic constraints on the backbone with a force constant of 10 kcal·mol*^−^*^1^·Å*^−^*^2^. This was followed by 40 ns of unrestrained equilibration using the NPT ensemble. All simulations were performed using the OpenMM7.7 biomolecular simulation package.^66^

#### 4.1.3 Production Simulations

Production MD was started using the equilibrated frames from each of the 18 systems. Once the first production runs were performed, frames for the next round of sampling were chosen using an adaptive sampling method. Details about adaptive sampling are explained in a separate section. Simulations were run on donated compute time provided by the Folding@Home distributed computing system. All simulations were run on OpenMM7.7. ^66^ Langevin Integrator was used to propagate the system and maintain the temperature of the system. System pressure was maintained at 1 bar using the MonteCarloMembraneBarostat used for maintaining uniform pressure in aniostropic membrane systems in OpenMM. Periodic boundary conditions were used. Particle Mesh Ewald Method (PME) ^67^ was used to compute contributions of long-range electrostatic interactions. The cutoff for switching to long-range interactions was set to 10 Å. Cutoff for non-bonded interactions was set to 10 Å. For each trajectory, frames were saved every 25000 steps, leading to a frame save rate of 0.1 ns. Total data collected for each system is reported in Table S1.

#### 4.1.4 Targeted MD simulations

To enable efficient sampling of the intermediate states during the activation process, the intermediate frames were also added to the list of starting points. These intermediate frames were sampled from the inactive to the active state, using a Targeted MD approach.^68^ The equilibrated inactive state was used as the input, and the equilibrated active state was set as the target. The overall transition simulation from the inactive to the active were implemented in NAMD^69^ and were performed over 1*µ*s. The intermediate frames were then used as seeds to start simulations, and combined with the rest of the starting points before the first round of adaptive sampling was performed. Overall, 5 replicates of the targeted MD simulations were performed for each of the 6 proteins.

### 4.2 Adaptive Sampling

The sampling process of the transition from the inactive to active state for a specific protein was further accelerated by using an adaptive sampling strategy.^44,45^ Several strategies have been proposed in the recent years,^70–75^ which use deep learning techniques to accelerate sampling over least counts adaptive sampling. Still, least count adaptive sampling performs on par with newer algorithms for exploratory sampling, while being computationally inexpensive.^45,72,74^ Simulations at end of each round were featurized based on C*α* distances between the conserved residues across the entire family (Tables S7-S12). The C*α* distance between a pair of residues pair was considered a feature if:

1. The collected data showed a difference of more than 4 Å between the highest and lowest values for that distance.
2. The maximum value of the distance was less than 8 Å.

The featurized trajectories were then clustered and simulation frames belonging to the least populated clusters were chosen as seeds for the next round of simulations. The following steps were followed for every specific protein:

1. Trajectories were seeded from the initial starting points.
2. The entire trajectory set was featurized using features that were able to differentiate between the inactive and the active states for the protein.
3. Clustering was performed on the featurized trajectories using k-means clustering.
4. Starting points for the next round of simulation were chosen from clusters with the least populations, to be run in parallel.
5. Dimensionality reduction was performed on the entire trajectory feature set using time-lagged Independent Component Analysis (tICA), to check whether the combined dataset for a protein had collectively sampled along the slowest kinetic process along the system (activation).

Details of the implementation of tICA are discussed in the next section. The adaptive strategy was iteratively performed on the entire trajectory set for a system, to generate multiple rounds of simulation data. Feature selection was done by first calculating the Residue-Residue Contact Score (RRCS) between the two starting points. Feature pairs (distances) that showed the ΔRRCS ≥ 3.5 were chosen as the features for characterizing the inactive to active transition for a particular protein. Feature sets used for each system are reported in Tables S7 - S12.

### 4.3 Dimensionality Reduction using tICA

time-lagged Independent Component Analysis^76^ is a dimensionality reduction technique that transforms the input time-series data (*ϕ_i_*) (MD simulation data in our case) by solving an eigenvalue problem. It finds a subspace of slowest processes in the system, making it ideal for an input to MSM construction. For a trajectory dataset, *χ_i_* and *χ_j_* define the trajectory feature sets, and *τ_tICA_* defines the tICA lagtime. tICA uses time-lagged (1) and instantaneous autocorrelation (2), and finds the slowest subspace **r***_i_* by solving the eigenvalue problem (3). The tICA lagtime *τ_tICA_* was chosen by first calculating the implied timescales *T* (where *T* = *^−^*^2/lnλ^). The value at which the *T* plateaued was chosen as *τ_tICA_* (30 ns for each protein system). Plots showing tIC1 v/s tIC2 (the two slowest processes captured by the system) are shown in Fig. S1.

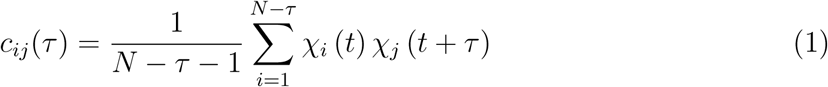

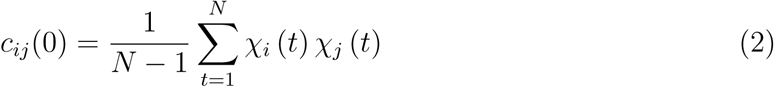

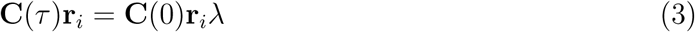

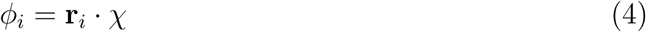

### 4.4 Markov State Model Construction

To recover the overall thermodynamics and kinetics of activation from short individual trajectories, a markov state model is constructed from the trajectory dataset. Markov state model is essentially a transition matrix **T**, where each element of the matrix *p*^^rev^(*τ*) describes the conditional probability of the system moving to state *j* at time *t* + *τ*, given that the system was in state *i* at time *t*.

Once tICA is performed, the slowest kinetic processes being modeled by the data are identified, the transformed data is clustered using k-means clustering to define the MSM microstates. Then, the transition probabilities between the MSM microstates are computed using the Maximum Likelihood Estimator,^77^ which maximizes the log probability -

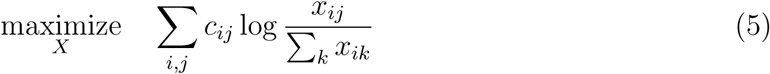

To solve the above optimization problem, a fixed-point iteration is performed using the following equation -

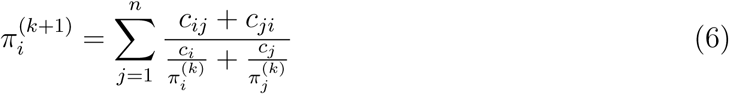

Once the *π_i_* values converge using the criteria ^1^*π*^(*k*+1)^ − *π*^(*k*)1^ *< ɛ*, the reversible transition probabilities are computed using -

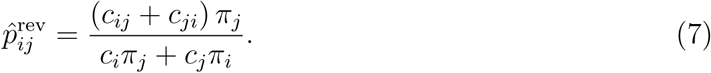

The optimum number of tIC components and number of microstates are MSM hyper-parameters, which were optimized using a grid-search approach. A proxy of the number of tIC components, the variational cutoff was used to account for the variance in the data. The combination of the variational cutoff and number of microstates which maximized the VAMP2 score was used for the construction of the final MSM (Fig. S5). To ensure that the model was maintaining Markovianity, the implied timescales were plotted against the MSM lagtime for each system, and a lagtime of 30 ns was chosen as the MSM lagtime, *τ_MSM_* (Fig. S6). The MSM was then validated using the Chapman-Kolmogorov test (Fig. S7-S12).

### 4.5 Trajectory Analysis

Trajectories were combined and stripped using cpptraj,^78^ a part of the AmberTools22 software suite. Scripts for analysis were written in python and are available on Github. mdtraj ^79^ was used for computing collective variables from trajectories. numpy, pandas, seaborn, matplotliband networkX python libraries were used for scientific computation. Trajectories were visualized using VMD,^80^ while ChimeraX ^81^ was used for preparing images.

#### 4.5.1 Construction of Free Energy Plots

Free Energy Plots were constructed by calculating the respective variable for the x and y axes, and 2D densities were computed using numpy’s histogram2d function. 200 bins were used for each dimension. Densities were weighed using the msm trajectory weights, and free energies were computed using the following formula -

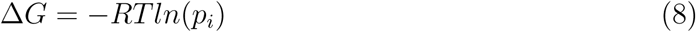

Errors for free energy plots were computed using bootstrapping - with 80% of the trajectories being iterated for 200 iterations.

#### 4.5.2 Kinetic Monte Carlo Simulations

Kinetic Monte Carlo Simulations were performed to identify molecular locks universal and unique to each protein’s activation. KMCs were simulated as a realization of the MSMs, using deeptime.markov.msm.MarkovStateModel.simulate. The microstate associated with the inactive conformation was provided as inputs to the function, and the realization was generated for 34,000 steps with timestep as the MSM lagtime *τ* (30 ns), thus totaling ∼1ms per trajectory,

## Acknowledgements

The authors would like to thank the Folding@Home distributed computing project for their aid with simulations. D.S. acknowledges support from NIH grant R35GM142745 and Cancer Center at Illinois. P.B. would like to thank A.T. Widiger Fellowship at the University of Illinois for their support. P.B. also thanks Caffe Paradiso at the University of Illinois for their support with providing an amenable space to conduct research and write manuscripts.

## 5 Data and Code Availability

Codes used for analysis are available at https://github.com/ShuklaGroup/ClassB1_Activation (GitHub). Trajectories and parameter files are available on Box: https://uofi.box.com/s/4g3xmumfmesb68y7tb0fn8wvhvycylrf

