## Supplementary Information for "All roads lead to Rome: Emergence of Mechanistic Divergence Shapes Activation of Class B1 G-Protein Coupled Receptors"

### List of Figures


S1 tICA plots . . . . . . . . . . . . . . . . . . . . . . . . . . . . . . . . . . . . . S3

S2 Differences between GCGR, GLP1R, PAC1R intermediate states . . . . . . . S4

S3 Differences between PTH1R, SCTR, CALCR intermediate states . . . . . . . S5

S4 Locks in SCTR Clade . . . . . . . . . . . . . . . . . . . . . . . . . . . . . . . S7

S5 MSM hyperparameter optimization . . . . . . . . . . . . . . . . . . . . . . . S8

S6 Implied Timescales Plot . . . . . . . . . . . . . . . . . . . . . . . . . . . . . S9

S7 Chapman-Kolmogorov test for GCGR MSM . . . . . . . . . . . . . . . . . . S10

S8 Chapman-Kolmogorov test for GLPR MSM . . . . . . . . . . . . . . . . . . . S11

S9 Chapman-Kolmogorov test for PTH1R MSM . . . . . . . . . . . . . . . . . . S12

S10 Chapman-Kolmogorov test for SCTR MSM . . . . . . . . . . . . . . . . . . . S13

S11 Chapman-Kolmogorov test for PAC1R MSM . . . . . . . . . . . . . . . . . . S14

S12 Chapman-Kolmogorov test for CALCR MSM . . . . . . . . . . . . . . . . . . S15


### List of Tables


S1 Simulated data per system . . . . . . . . . . . . . . . . . . . . . . . . . . . S16

S2 PDB IDs of the systems simulated in this study . . . . . . . . . . . . . . . . S17

S3 Mutations performed to revert to wild type . . . . . . . . . . . . . . . . . . . S18

S4 Modelled Residues in each system . . . . . . . . . . . . . . . . . . . . . . . . S19

S5 Protonations applied to acidic residues in the systems simulated in this study S23

S6 System sizes of the systems simulated in this study . . . . . . . . . . . . . . S24

S7 Adaptive Sampling Metrics for GCGR . . . . . . . . . . . . . . . . . . . . . S25

S8 Adaptive Sampling Metrics for GLP1R . . . . . . . . . . . . . . . . . . . . . S26

S9 Adaptive Sampling Metrics for PTH1R . . . . . . . . . . . . . . . . . . . . . S27

S10 Adaptive Sampling Metrics for SCTR . . . . . . . . . . . . . . . . . . . . . . S28

S11 Adaptive Sampling Metrics for PAC1R . . . . . . . . . . . . . . . . . . . . . S29

S12 Adaptive Sampling Metrics for CALCR . . . . . . . . . . . . . . . . . . . . . . S30

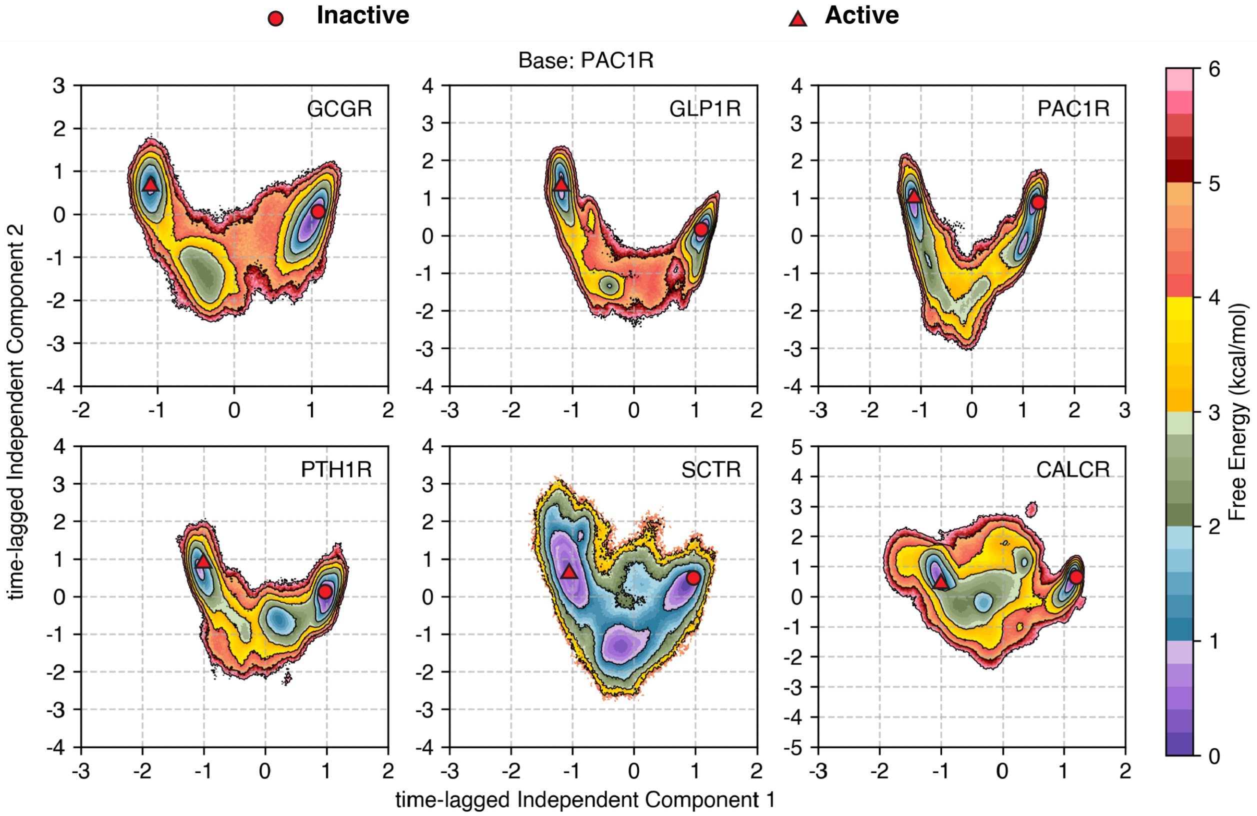


Supplementary Figure S1: Projection of the first two time-lagged Independent Component Projections for each system in this study. As evidenced by the plot, the first tIC component along the x axis, shows separation for the active and inactive states for all 6 proteins, showing that the slowest process captured by the overall dataset for each protein is activation.

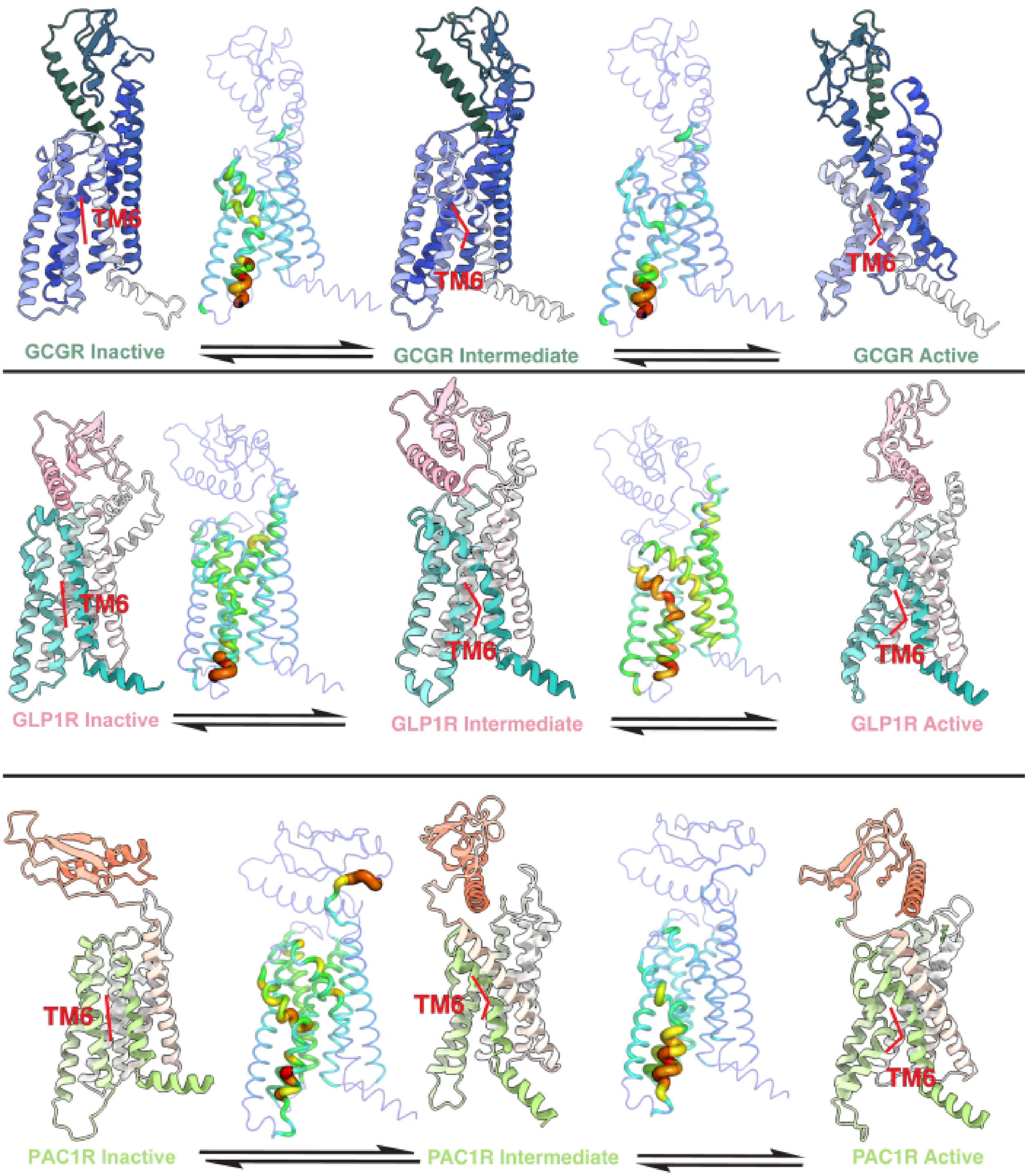


Supplementary Figure S2: Differences between GCGR, GLP1R, PAC1R intermediate states. For each protein, the inactive, intermediate, and active states are shown. Schematics above the arrows show the transitions that happen between corresponding states. The transitions between any two states are colored according to the symmetric KL-Divergence between the start and the end state, with blue being the lowest and red being the highest. States correspond to (a) GCGR inactive (b) GCGR intermediate (c) GCGR active (d) GLP1R inactive (e) GLP1R intermediate (f) GLP1R active (g) PAC1R inactive (h) PAC1R intermediate (i) PAC1R active states.

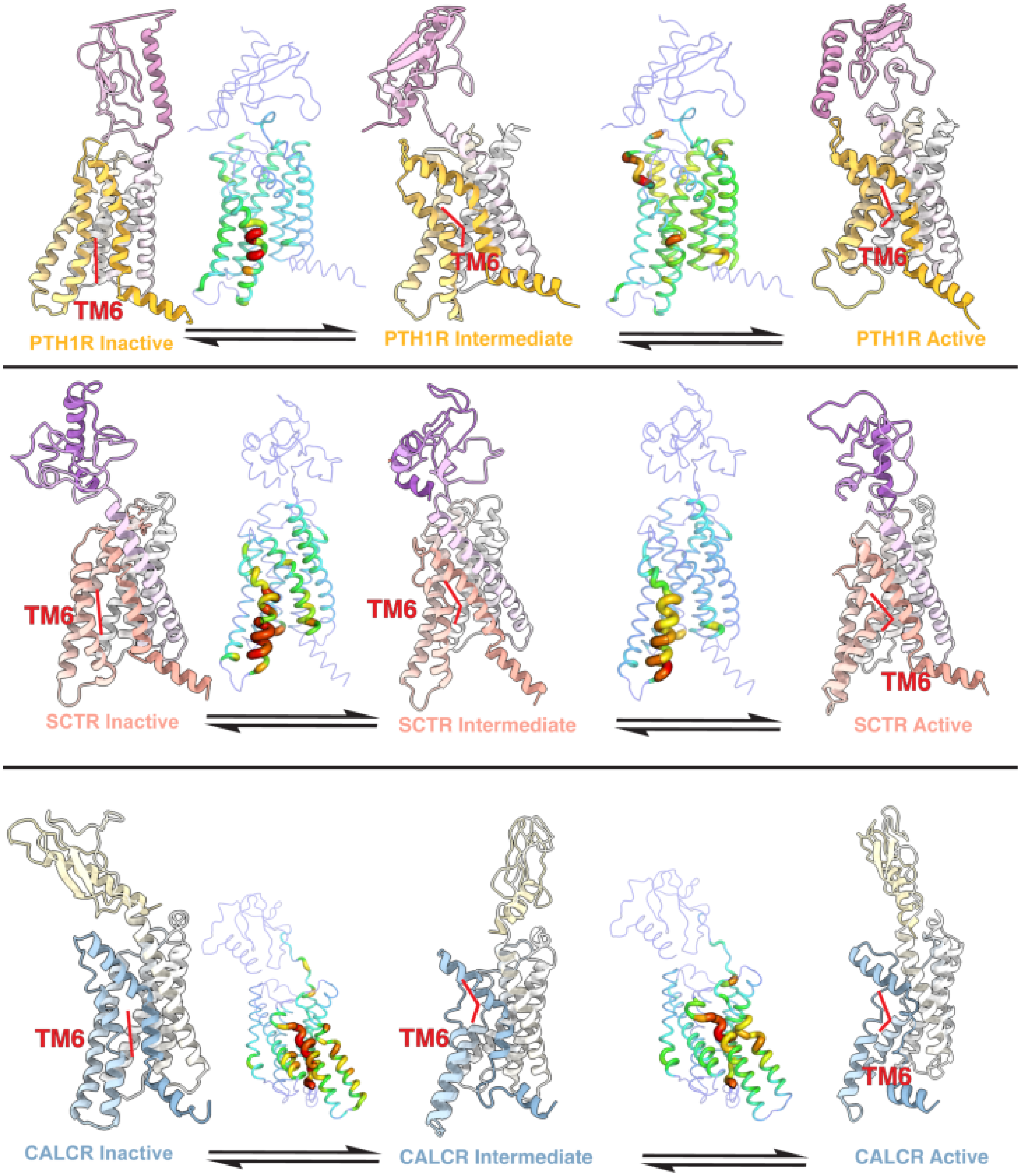


Supplementary Figure S3: Differences between PTH1R, SCTR, CALCR intermediate states. For each protein, the inactive, intermediate, and active states are shown. Schematics above the arrows show the transitions that happen between corresponding states. The transitions between any two states are colored according to the symmetric KL-Divergence between the start and the end state, with blue being the lowest and red being the highest. States correspond to (a) PTH1R inactive (b) PTH1R intermediate (c) PTH1R active (d) SCTR inactive (e) SCTR intermediate (f) SCTR active (g) CALCR inactive (h) CALCR intermediate (i) CALCR active states.

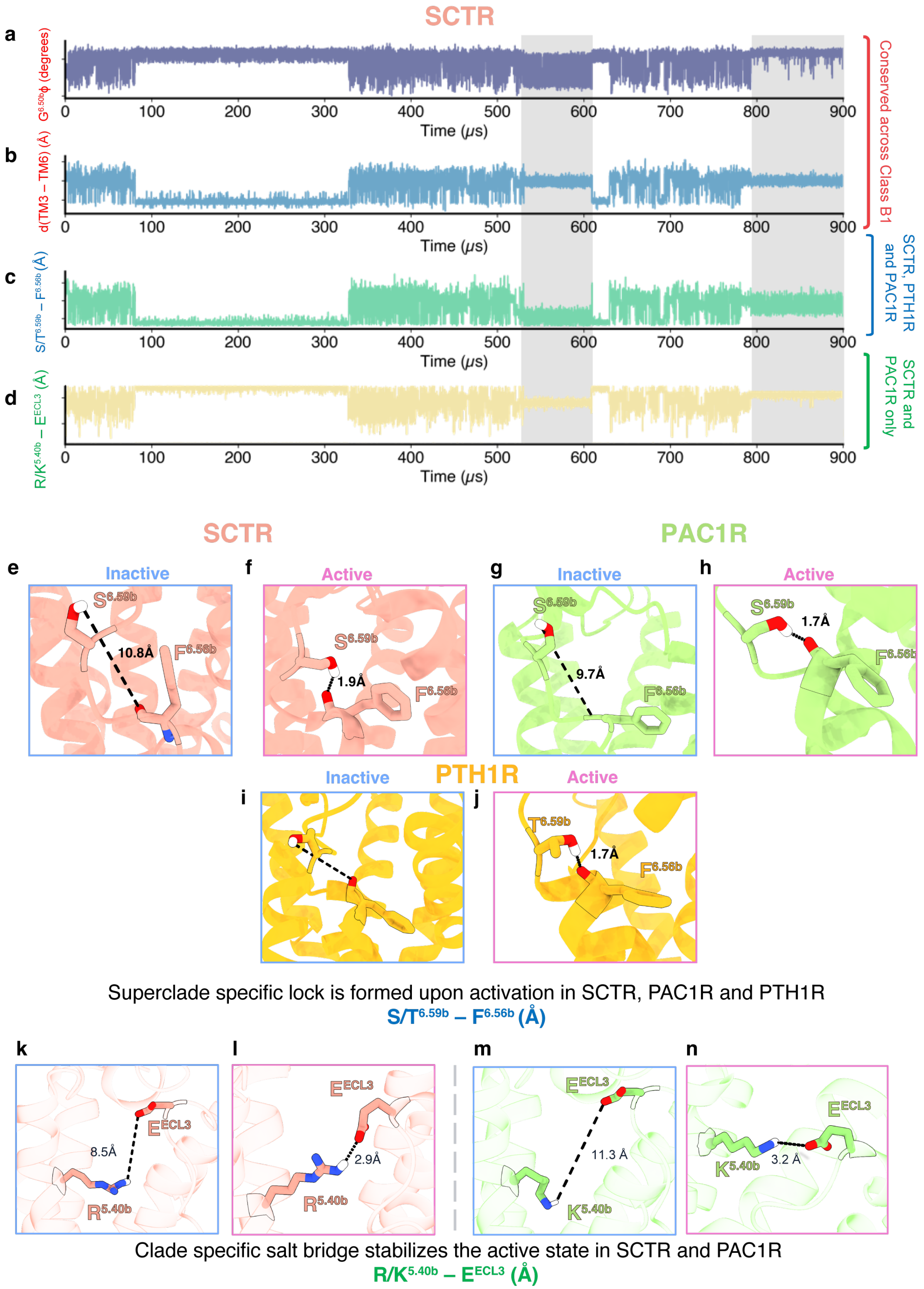
SCTR
a
G6.50b φ (degrees)
b
d(TM3 – TM6) (Å)
c
S/T6.59b – F6.56b (Å)
d
R/K5.40b – EECL3 (Å)
0
100
200
300
400
500
600
700
800
900
Time (μs)
Conserved across Class B1
SCTR, PTH1R and PAC1R
SCTR and PAC1R only
SCTR
PAC1R
e
Inactive
S6.59b
10.8Å
F6.56b
f
Active
S6.59b
1.9Å
F6.56b
g
Inactive
S6.59b
9.7Å
F6.56b
h
Active
S6.59b
1.7Å
F6.56b
PTH1R
i
Inactive
j
Active
T6.59b
1.7Å
F6.56b
Superclade specific lock is formed upon activation in SCTR, PAC1R and PTH1R
S/T6.59b – F6.56b (Å)
k
8.5Å
EECL3
R5.40b
l
EECL3
2.9Å
R5.40b
m
EECL3
11.3 Å
K5.40b
n
EECL3
K5.40b
3.2 Å
Clade specific salt bridge stabilizes the active state in SCTR and PAC1R
R/K5.40b – EECL3 (Å)

Supplementary Figure S4: Clade-wise activation mechanism in the SCTR-PTH1R-PAC1R clade. (a-e) KMC simulations performed for SCTR. Active confirmations are highlighted in light grey. The plots show (a) backbone dihedral angle $\phi$ at $G^{6.50b}$, (b) distance between TM3 and TM6 measured at $L^{3.57b}$ and $T^{6.42b}$, (c) distance between $S/T^{6.59b}$ and $F^{6.56b}$ (d) distance between $R/K^{5.40b}$ and $E^{ECL3}$ (e-j) hydrogen bonds between $S/T^{6.59b}$ and $F^{6.56b}$ form as the proteins transition from inactive (e,g,i) and active (f,h,j), for SCTR (e,f), PAC1R (g,h) and PTH1R (i,j). (k-n) salt bridges between $R/K^{5.40b}$ and $E^{ECL3}$ form as the proteins transition from inactive (k,m) and active (l,n), for SCTR (k,l) and PAC1R (m, n).

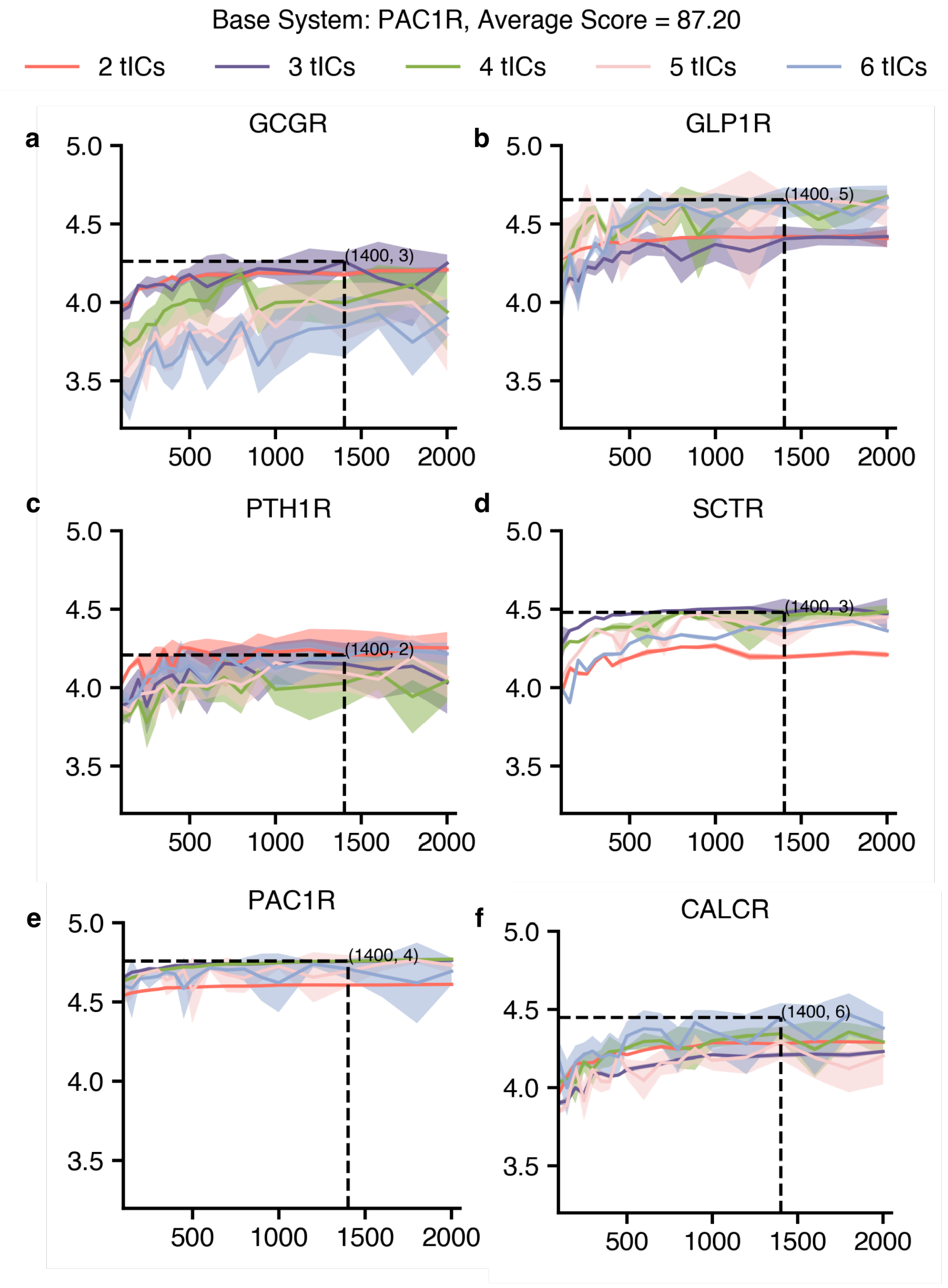


Supplementary Figure S5: VAMPscores v/s number of clusters for each system. Scores were calculated from 100-1000 clusters and at 4 variational cutoffs - 50%, 65%, 80% and 95% , each corresponding to different number of tICs considered while calculating the score.

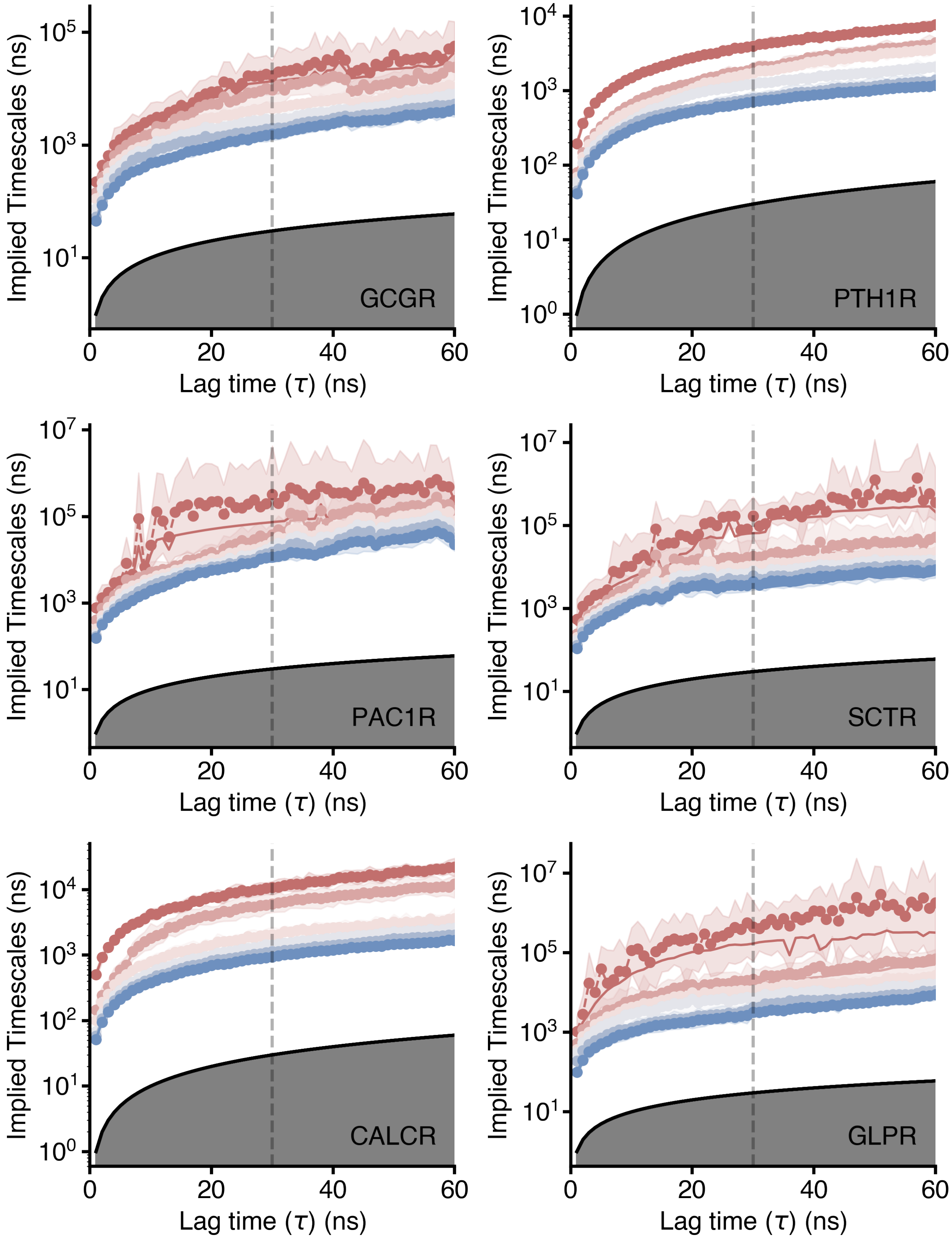


Supplementary Figure S6: Implied Timescales v/s MSM lagtime for the 6 proteins considered in this study. A lagtime of 30ns (dotted vertical line) was chosen for each MSM.

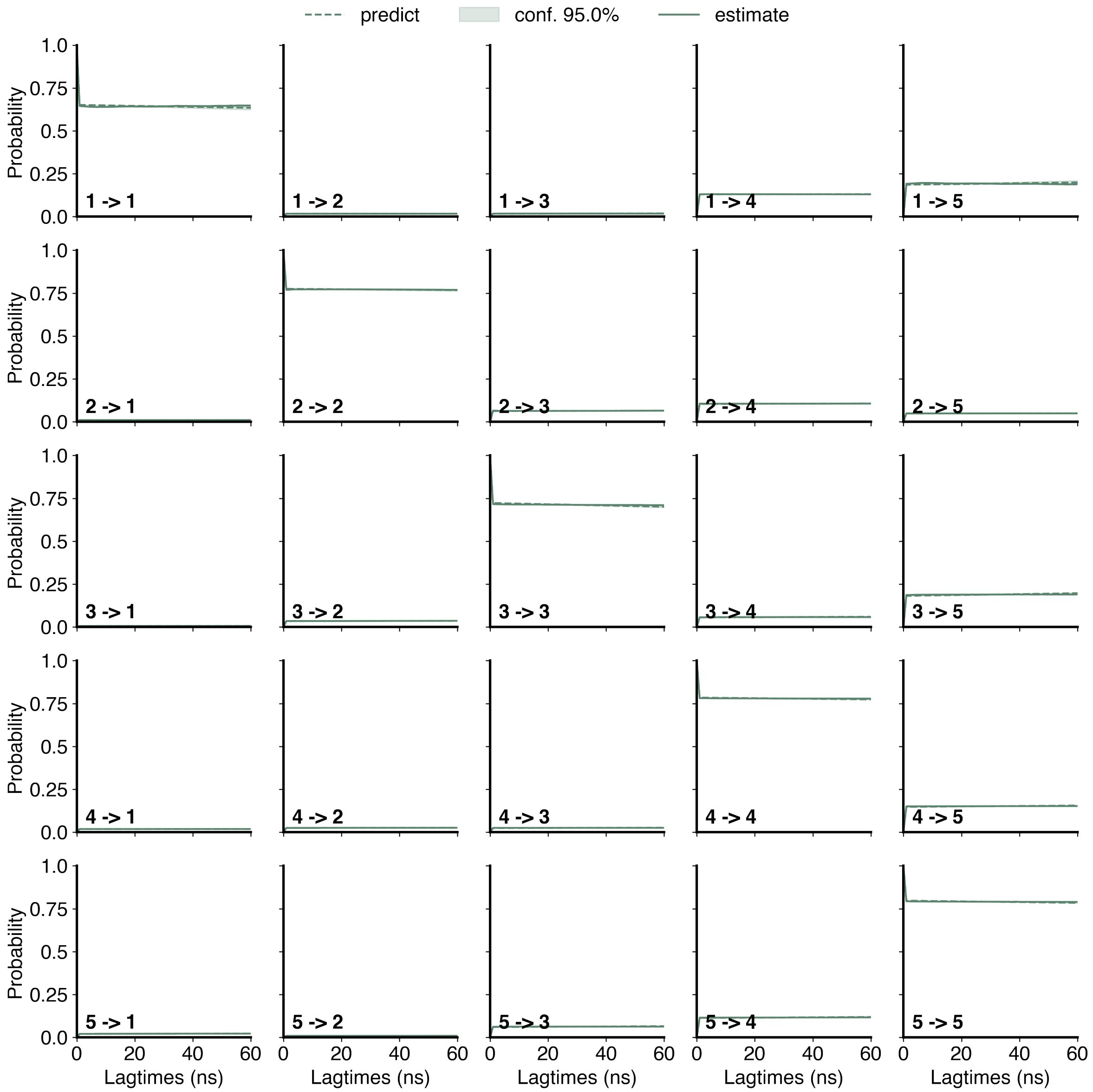


Supplementary Figure S7: Chapman Kolmogorov Test for the MSM built using GCGR Apo data. 5 macrostates were considered.

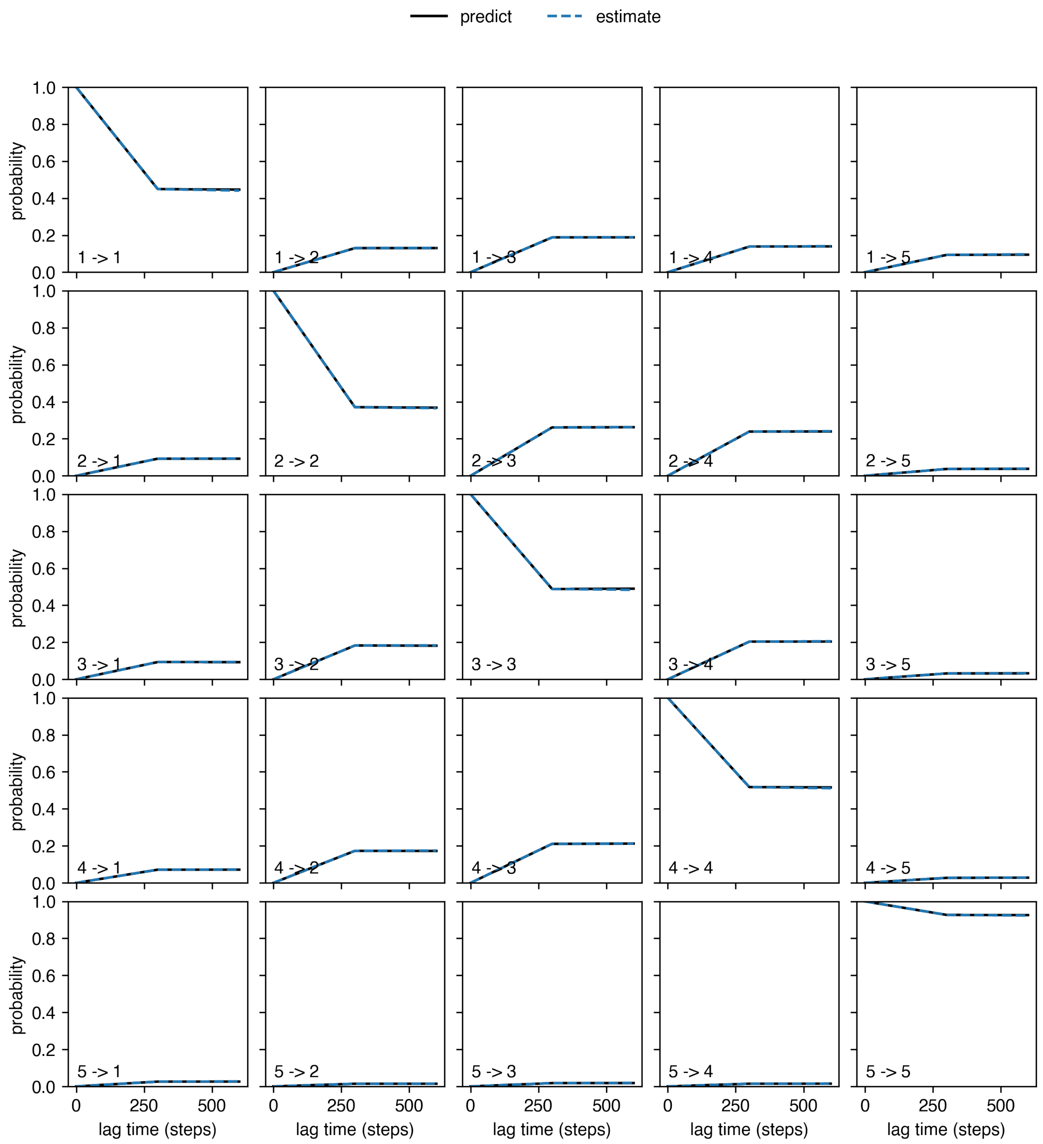


Supplementary Figure S8: Chapman Kolmogorov Test for the MSM built using GLPR Apo data. 5 macrostates were considered.

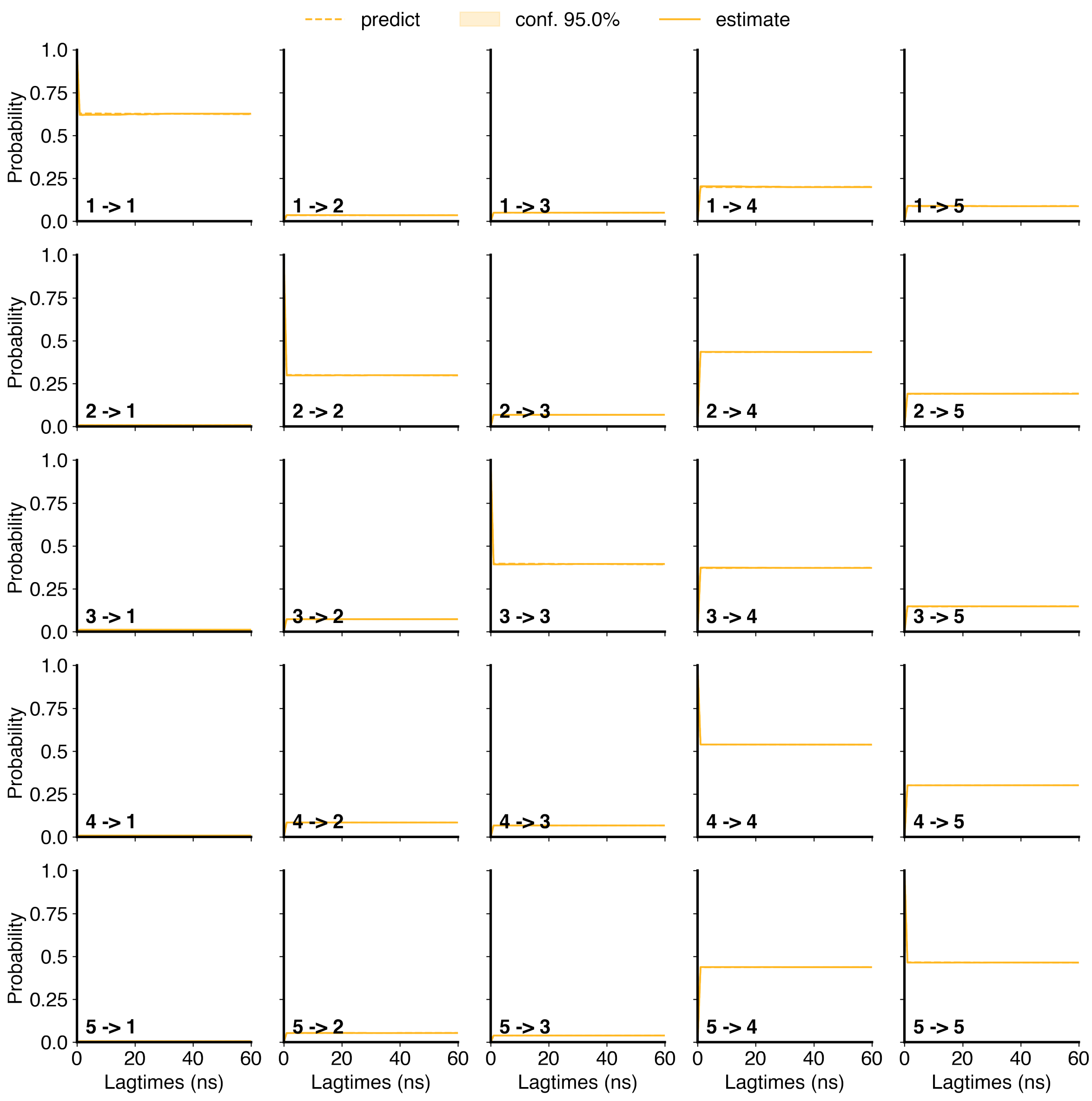


Supplementary Figure S9: Chapman Kolmogorov Test for the MSM built using PTH1R Apo data. 5 macrostates were considered.

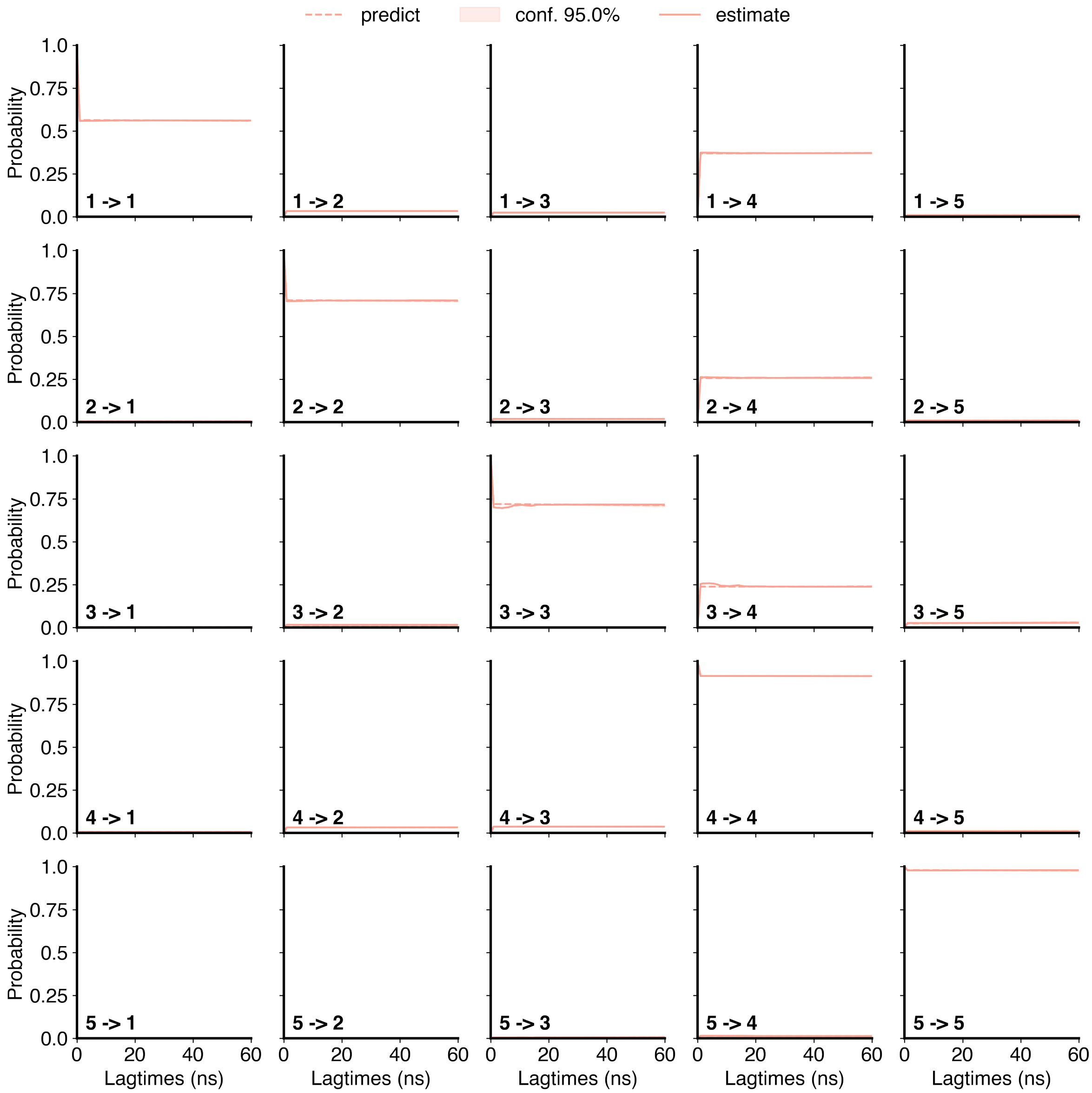


Supplementary Figure S10: Chapman Kolmogorov Test for the MSM built using SCTR Apo data. 5 macrostates were considered.

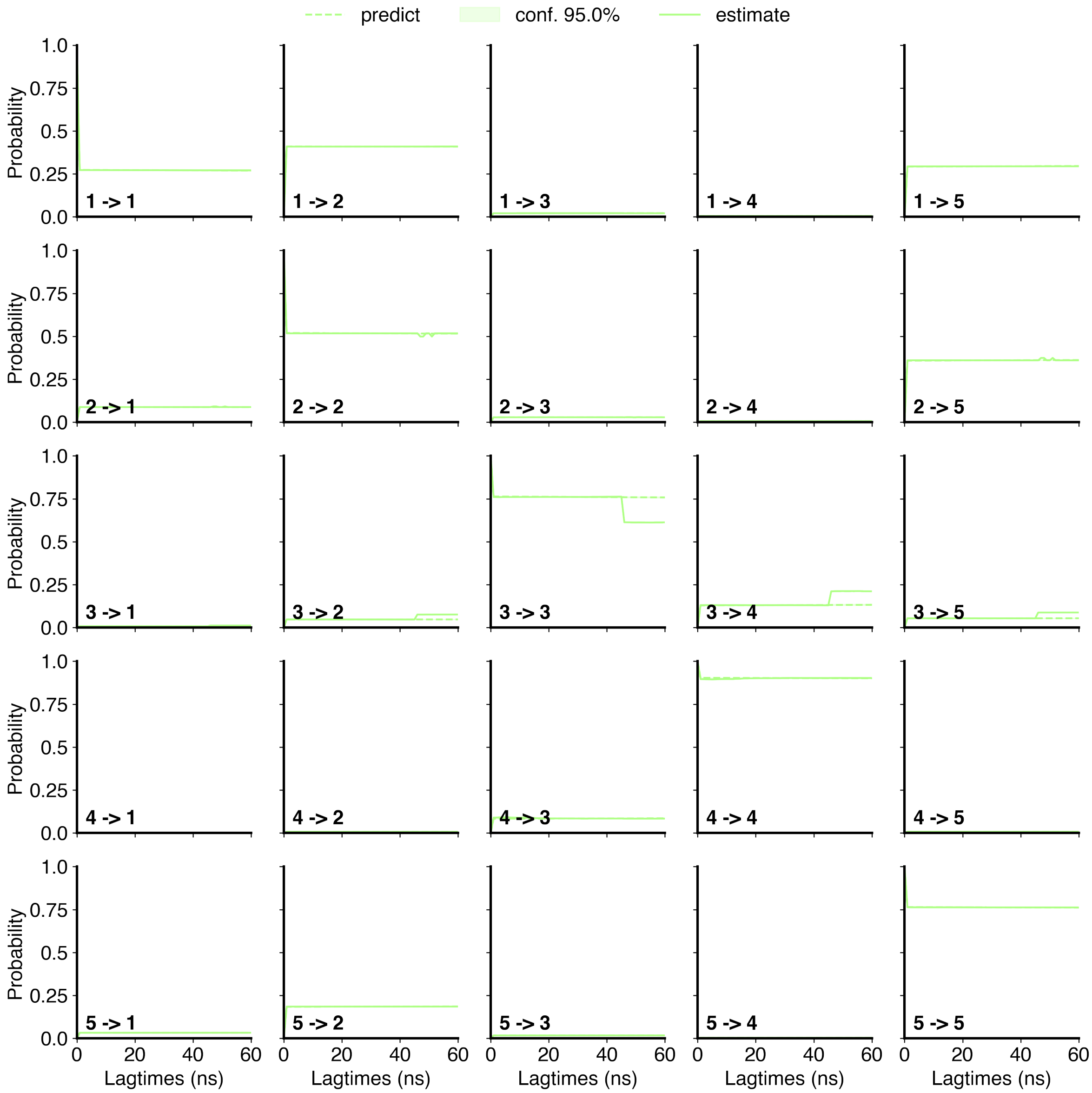


Supplementary Figure S11: Chapman Kolmogorov Test for the MSM built using PAC1R Apo data. 5 macrostates were considered.

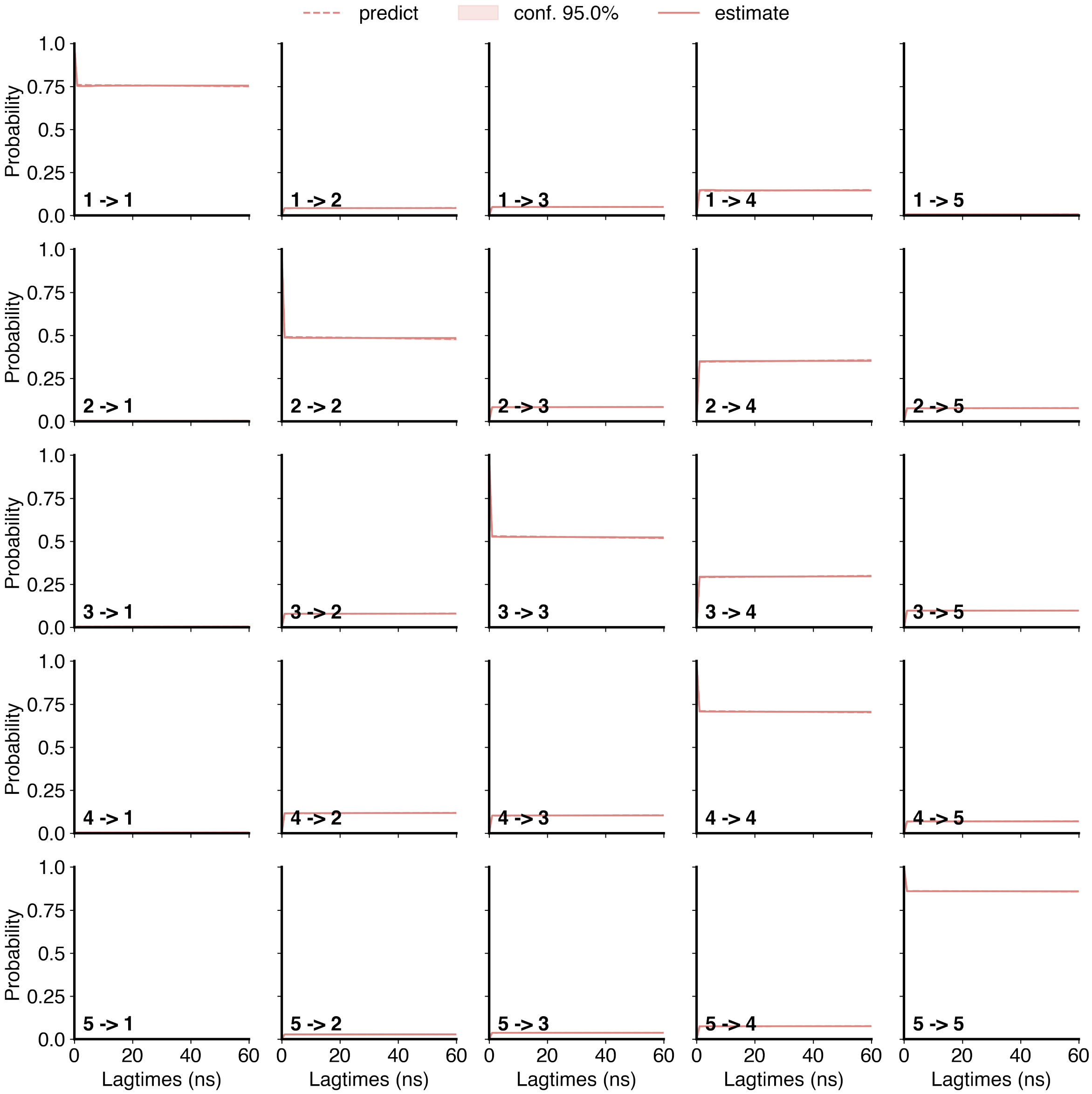


Supplementary Figure S12: Chapman Kolmogorov Test for the MSM built using CALCR Apo data. 5 macrostates were considered.

**Supplementary Table S1: Amount of simulations performed for each system.**

| Protein | State | Conformation | Total Data |
|---|---|---|---|
| GCGR | APO | Inactive | 560 $\mu$s |
| | APO | Active | 360 $\mu$s |
| | Holo | | 50 $\mu$s |
| GLP1R | APO | Inactive | 700 $\mu$s |
| | APO | Active | 592 $\mu$s |
| | Holo | | 50 $\mu$s |
| SCTR | APO | Inactive | 652 $\mu$s |
| | APO | Active | 260 $\mu$s |
| | Holo | | 50 $\mu$s |
| PAC1R | APO | Inactive | 220 $\mu$s |
| | APO | Active | 180 $\mu$s |
| | Holo | | 50 $\mu$s |
| PTH1R | APO | Inactive | 800 $\mu$s |
| | APO | Active | 124 $\mu$s |
| | Holo | | 50 $\mu$s |
| CALCR | APO | Inactive | 400 $\mu$s |
| | APO | Active | 114 $\mu$s |
| | Holo | | 50 $\mu$s |
| **Total simulation time** | | | 5262 $\mu$s |

**Supplementary Table S2: PDB IDs info of the systems simulated in this study.**

| Protein | State | Conformation | PDB ID | Reference |
|---|---|---|---|---|
| GCGR | APO | Inactive | 5YQZ | Zhang et al.[1] |
| | APO<br>HOLO | Active | 6WPW | Hilger et al.[2] |
| GLP1R | APO | Inactive | 6LN2 | Wu et al.[3] |
| | APO<br>HOLO | Active | 6X18 | Zhang et al.[4] |
| PTH1R | APO | Inactive | 6FJ3 | Ehrenmann et al.[5] |
| | APO<br>HOLO | Active | 6NBF | Zhao et al.[6] |
| SCTR | APO | Inactive | Alpha-fold2 Multistate Model | Heo and Feig[7][8] |
| | APO<br>HOLO | Active | 6WZG | Dong et al.[9] |
| PAC1R | APO | Inactive | Alpha-fold2 Multistate Model | Heo and Feig[7][10] |
| | APO<br>HOLO | Active | 6M1I | Wang et al.[11] |
| CALCR | APO | Inactive | Alpha-fold2 Multistate Model | Heo and Feig[7][12] |
| | APO<br>HOLO | Active | 7TYO | Cao et al.[13] |

**Supplementary Table S3: Mutations performed to revert to wild type**

| Protein | PDB ID | Mutation/s |
|---|---|---|
| GCGR | 5YQZ | A173R |
| | 6WPW | - |
| GLP1R | 6LN2 | C193S, F196I, A225S, C233M, A271S, C317I, I318G, A346K, F347C, C361G, D387E |
| | 6X18 | - |
| PTH1R | 6FJ3 | C191Y, M240K, A300L, K312M, I334V, N359K, A407L, L426A, R440Q, A458I |
| | 6NBF | A188G |
| SCTR | 6WZG | - |
| PAC1R | 6M1I | - |
| CALCR | 7TYO | A366S |

**Supplementary Table S4: Modelled Residues in each system**

| Protein | PDB ID | Modelled Residues | Restraints added | Location |
|---|---|---|---|---|
| GCGR | 5YQZ | L258 | None | ICL2 |
| | | P259 | None | ICL2 |
| | 6WPW | G109 | None | ECD |
| | | P110 | None | ECD |
| | | D111 | None | ECD |
| | | W433 | $\alpha$-helical | Helix-8 |
| | | E434 | $\alpha$-helical | Helix-8 |
| GLP1R | 6LN2 | S129 | None | ECD |
| | | K130 | None | ECD |
| | | R131 | None | ECD |
| | | G132 | None | ECD |
| | | E133 | None | ECD |
| | | R134 | None | ECD |
| | | S258 | None | ICL2 |
| | | V259 | None | ICL2 |
| | | L260 | None | ICL2 |
| | | S261 | None | ICL2 |
| | 6X18 | K130 | None | ECD |
| | | R131 | None | ECD |
| | | G132 | None | ECD |
| | | E133 | None | ECD |
| | | R134 | None | ECD |
| | | S135 | None | ECD |
| | | L339 | None | ICL3 |

Continued on next page

| Protein | PDB ID | Modelled Residues | Restraints added | Location |
|---|---|---|---|---|
| | | M340 | None | ICL3 |
| | | C341 | None | ICL3 |
| | | K342 | None | ICL3 |
| | | T343 | None | ICL3 |
| PTH1R | 6FJ3 | L389 | None | ICL2 |
| | | R390 | None | ICL2 |
| | | E391 | None | ICL2 |
| | | T392 | None | ICL2 |
| | | N393 | None | ICL2 |
| | | A394 | None | ICL2 |
| | | G395 | None | ICL2 |
| | | R396 | None | ICL2 |
| | | C397 | None | ICL2 |
| | 6NBF | L56 | None | ECD |
| | | Q57 | None | ECD |
| | | R58 | None | ECD |
| | | P59 | None | ECD |
| | | A60 | None | ECD |
| | | A394 | None | ICL2 |
| | | G395 | None | ICL2 |
| | | R396 | None | ICL2 |
| | | C397 | None | ICL2 |
| | | D398 | None | ICL2 |
| SCTR | 6WZG | - | - | - |

Continued on next page

| Protein | PDB ID | Modelled Residues | Restraints added | Location |
|---|---|---|---|---|
| PAC1R | 6M1I | N42 | None | ECD |
| | | E43 | None | ECD |
| | | L44 | None | ECD |
| | | M45 | None | ECD |
| | | G46 | None | ECD |
| | | F47 | None | ECD |
| | | N48 | None | ECD |
| | | D49 | None | ECD |
| | | S50 | None | ECD |
| | | S51 | None | ECD |
| | | E120 | None | ECD |
| | | D121 | None | ECD |
| | | Y139 | None | ECD |
| | | E140 | None | ECD |
| | | S141 | None | ECD |
| | | E142 | None | ECD |
| | | T143 | None | ECD |
| | | M340 | None | ICL3 |
| | | G341 | None | ICL3 |
| | | G342 | None | ICL3 |
| | | N343 | None | ICL3 |
| | | E344 | None | ICL3 |
| | | S345 | None | ICL3 |
| | | Q61 | None | ECD |
| | | L62 | None | ECD |

Continued on next page

| Protein | PDB ID | Modelled Residues | Restraints added | Location |
| --- | --- | --- | --- | --- |
| | | P63 | None | ECD |
| | | A64 | None | ECD |
| | | Y65 | None | ECD |
| | | Q66 | None | ECD |
| | | G67 | None | ECD |
| | | E68 | None | ECD |
| | | G69 | None | ECD |
| | | P70 | None | ECD |
| | | E114 | None | ECD |
| | | K115 | None | ECD |
| | | G116 | None | ECD |

**Supplementary Table S5: Protonations applied to acidic residues in the systems simulated in this study.**

| Protein | Protonation/s |
|---|---|
| GCGR | H361, H372 |
| GLP1R | H99, E128, E247 |
| PTH1R | H114, H143, H223 |
| SCTR | - |
| PAC1R | H68, H129, H189, H218 |
| CALCR | H336 |

**Supplementary Table S6: System sizes of the systems simulated in this study.**

| Protein | Conformation | State | System Size $(\AA^3)$ | Atoms |
|---|---|---|---|---|
| GCGR | Inactive | APO | $89.4 \times 89.4 \times 144.7$ | 107,187 |
| | Active | APO | $90.1 \times 90.1 \times 142.9$ | 107,715 |
| | | HOLO | $89.8 \times 89.8 \times 142.8$ | 107,962 |
| GLP1R | Inactive | APO | $90.2 \times 90.2 \times 138.7$ | 106,917 |
| | Active | APO | $89.5 \times 89.5 \times 143.7$ | 107,489 |
| | | HOLO | $89.8 \times 89.8 \times 143.9$ | 107,708 |
| PTH1R | Inactive | APO | $89.1 \times 89.1 \times 151.2$ | 111,616 |
| | Active | APO | $88.8 \times 88.8 \times 151.0$ | 111,405 |
| | | HOLO | $89.4 \times 89.4 \times 151.2$ | 111,974 |
| SCTR | Inactive | APO | $90.1 \times 90.1 \times 145.6$ | 109,586 |
| | Active | APO | $90.1 \times 90.1 \times 145.3$ | 109,533 |
| | | HOLO | $89.9 \times 89.9 \times 145.8$ | 110,266 |
| PAC1R | Inactive | APO | $89.8 \times 89.8 \times 136.2$ | 100,960 |
| | Active | APO | $89.9 \times 89.9 \times 136.1$ | 101,265 |
| | | HOLO | $89.9 \times 89.9 \times 135.9$ | 101,757 |
| CALCR | Inactive | APO | $89.7 \times 89.7 \times 122.1$ | 90,355 |
| | Active | APO | $89.2 \times 89.2 \times 121.3$ | 89,975 |
| | | HOLO | $89.2 \times 89.2 \times 121.5$ | 90,538 |

| Adaptive Sampling Metrics - GCGR | | | | | | | | | | | |
|---|---|---|---|---|---|---|---|---|---|---|---|
| Residue 1 | W.N. 1 | Residue 2 | W.N. 2 | Residue 1 | W.N. 1 | Residue 2 | W.N. 2 | Residue 1 | W.N. 1 | Residue 2 | W.N. 2 |
| C58 | ECD | W68 | ECD | C224 | 3.29 | W295 | ECL2 | L354 | 6.45 | L357 | 6.48 |
| C58 | ECD | C100 | ECD | N238 | 3.43 | W241 | 3.46 | L354 | 6.45 | G359 | 6.50 |
| C58 | ECD | G104 | ECD | N238 | 3.43 | G271 | 4.49 | L354 | 6.45 | Q392 | 7.49 |
| D63 | ECD | C67 | ECD | N238 | 3.43 | W272 | 3.50 | L354 | 6.45 | V396 | 7.53 |
| D63 | ECD | W68 | ECD | N238 | 3.43 | L357 | 6.48 | P356 | 6.47 | L357 | 6.48 |
| C67 | ECD | W68 | ECD | N238 | 3.43 | G359 | 6.50 | P356 | 6.47 | G359 | 6.50 |
| C67 | ECD | C81 | ECD | W241 | 3.46 | E245 | 3.50 | P356 | 6.47 | F391 | 7.48 |
| C67 | ECD | P82 | ECD | W241 | 3.46 | G271 | 4.49 | P356 | 6.47 | Q392 | 7.49 |
| W68 | ECD | P82 | ECD | W241 | 3.46 | W272 | 3.50 | P356 | 6.47 | G393 | 7.50 |
| C81 | ECD | P82 | ECD | E245 | 3.50 | Y248 | 3.53 | P356 | 6.47 | V396 | 7.53 |
| C100 | ECD | G104 | ECD | E245 | 3.50 | L249 | 3.54 | L357 | 6.48 | G359 | 6.50 |
| G148 | 1.46 | S152 | 1.50 | E245 | 3.50 | N318 | 5.50 | L357 | 6.48 | F391 | 7.48 |
| G148 | 1.46 | F391 | 7.48 | E245 | 3.50 | T351 | 6.42 | L357 | 6.48 | Q392 | 7.49 |
| G148 | 1.46 | Q392 | 7.49 | E245 | 3.50 | L354 | 6.45 | L357 | 6.48 | G393 | 7.50 |
| G148 | 1.46 | G393 | 7.50 | E245 | 3.50 | L357 | 6.48 | L357 | 6.48 | V396 | 7.53 |
| S152 | 1.50 | L156 | 1.54 | E245 | 3.50 | G359 | 6.50 | G359 | 6.50 | F391 | 7.48 |
| S152 | 1.50 | F391 | 7.48 | Y248 | 3.53 | L249 | 3.54 | G359 | 6.50 | Q392 | 7.49 |
| S152 | 1.50 | Q392 | 7.49 | L249 | 3.54 | N318 | 5.50 | G359 | 6.50 | G393 | 7.50 |
| S152 | 1.50 | G393 | 7.50 | L249 | 3.54 | T351 | 6.42 | G359 | 6.50 | V396 | 7.53 |
| S152 | 1.50 | V396 | 7.53 | L249 | 3.54 | L354 | 6.45 | F391 | 7.48 | Q392 | 7.49 |
| L156 | 1.54 | G393 | 7.50 | L249 | 3.54 | L357 | 6.48 | F391 | 7.48 | G393 | 7.50 |
| L156 | 1.54 | V396 | 7.53 | G271 | 4.49 | W272 | 3.50 | F391 | 7.48 | V396 | 7.53 |
| L156 | 1.54 | C401 | 7.58 | G271 | 4.49 | P275 | 4.53 | Q392 | 7.49 | G393 | 7.50 |
| C171 | 2.44 | R173 | 2.46 | W272 | 3.50 | P275 | 4.53 | Q392 | 7.49 | V396 | 7.53 |
| C171 | 2.44 | E406 | 8.49 | P275 | 4.53 | P310 | 5.42 | G393 | 7.50 | V396 | 7.53 |
| C171 | 2.44 | V407 | 8.50 | C294 | ECL2 | W295 | ECL2 | V396 | 7.53 | C401 | 7.58 |
| R173 | 2.46 | H177 | 2.50 | N318 | 5.50 | T351 | 6.42 | C401 | 7.58 | F402 | 7.59 |
| R173 | 2.46 | E245 | 3.50 | N318 | 5.50 | L354 | 6.45 | C401 | 7.58 | N404 | 8.47 |
| R173 | 2.46 | Y248 | 3.53 | N318 | 5.50 | P356 | 6.47 | C401 | 7.58 | E406 | 8.49 |
| R173 | 2.46 | L249 | 3.54 | N318 | 5.50 | L357 | 6.48 | C401 | 7.58 | V407 | 8.50 |
| R173 | 2.46 | E406 | 8.49 | N318 | 5.50 | G359 | 6.50 | F402 | 7.59 | N404 | 8.47 |
| H177 | 2.50 | W241 | 3.46 | T351 | 6.42 | L354 | 6.45 | F402 | 7.59 | V407 | 8.50 |
| H177 | 2.50 | E245 | 3.50 | T351 | 6.42 | P356 | 6.47 | N404 | 8.47 | E406 | 8.49 |
| H177 | 2.50 | V396 | 7.53 | T351 | 6.42 | G359 | 6.50 | N404 | 8.47 | V407 | 8.50 |
| H177 | 2.50 | C401 | 7.58 | L354 | 6.45 | P356 | 6.47 | E406 | 8.49 | V407 | 8.50 |
| C224 | 3.29 | C294 | ECL2 | | | | | | | | |

**Supplementary Table S7: Adaptive Sampling Metrics for GCGR. Residue contacts between Residue 1 and Residue 2 were used as metrics for Adaptive Sampling. The Wootten numbering for each residue is reported as well.**

| Adaptive Sampling Metrics - GLP1R | | | | | | | | | | | |
|---|---|---|---|---|---|---|---|---|---|---|---|
| Residue 1 | W.N. 1 | Residue 2 | W.N. 2 | Residue 1 | W.N. 1 | Residue 2 | W.N. 2 | Residue 1 | W.N. 1 | Residue 2 | W.N. 2 |
| C62 | ECD | W72 | ECD | C226 | 3.29 | W297 | ECL2 | L356 | 6.45 | L359 | 6.48 |
| C62 | ECD | C104 | ECD | N240 | 3.43 | W243 | 3.46 | L356 | 6.45 | G361 | 6.50 |
| C62 | ECD | G108 | ECD | N240 | 3.43 | G273 | 4.49 | L356 | 6.45 | Q394 | 7.49 |
| D67 | ECD | C71 | ECD | N240 | 3.43 | W274 | 3.50 | L356 | 6.45 | V398 | 7.53 |
| D67 | ECD | W72 | ECD | N240 | 3.43 | L359 | 6.48 | P358 | 6.47 | L359 | 6.48 |
| C71 | ECD | W72 | ECD | N240 | 3.43 | G361 | 6.50 | P358 | 6.47 | G361 | 6.50 |
| C71 | ECD | C85 | ECD | W243 | 3.46 | E247 | 3.50 | P358 | 6.47 | F393 | 7.48 |
| C71 | ECD | P86 | ECD | W243 | 3.46 | G273 | 4.49 | P358 | 6.47 | Q394 | 7.49 |
| W72 | ECD | P86 | ECD | W243 | 3.46 | W274 | 3.50 | P358 | 6.47 | G395 | 7.50 |
| C85 | ECD | P86 | ECD | E247 | 3.50 | Y250 | 3.53 | P358 | 6.47 | V398 | 7.53 |
| C104 | ECD | G108 | ECD | E247 | 3.50 | L251 | 3.54 | L359 | 6.48 | G361 | 6.50 |
| G151 | 1.46 | S155 | 1.50 | E247 | 3.50 | N320 | 5.50 | L359 | 6.48 | F393 | 7.48 |
| G151 | 1.46 | F393 | 7.48 | E247 | 3.50 | T353 | 6.42 | L359 | 6.48 | Q394 | 7.49 |
| G151 | 1.46 | Q394 | 7.49 | E247 | 3.50 | L356 | 6.45 | L359 | 6.48 | G395 | 7.50 |
| G151 | 1.46 | G395 | 7.50 | E247 | 3.50 | L359 | 6.48 | L359 | 6.48 | V398 | 7.53 |
| S155 | 1.50 | L159 | 1.54 | E247 | 3.50 | G361 | 6.50 | G361 | 6.50 | F393 | 7.48 |
| S155 | 1.50 | F393 | 7.48 | Y250 | 3.53 | L251 | 3.54 | G361 | 6.50 | Q394 | 7.49 |
| S155 | 1.50 | Q394 | 7.49 | L251 | 3.54 | N320 | 5.50 | G361 | 6.50 | G395 | 7.50 |
| S155 | 1.50 | G395 | 7.50 | L251 | 3.54 | T353 | 6.42 | G361 | 6.50 | V398 | 7.53 |
| S155 | 1.50 | V398 | 7.53 | L251 | 3.54 | L356 | 6.45 | F393 | 7.48 | Q394 | 7.49 |
| L159 | 1.54 | G395 | 7.50 | L251 | 3.54 | L359 | 6.48 | F393 | 7.48 | G395 | 7.50 |
| L159 | 1.54 | V398 | 7.53 | G273 | 4.49 | W274 | 3.50 | F393 | 7.48 | V398 | 7.53 |
| L159 | 1.54 | C403 | 7.58 | G273 | 4.49 | P277 | 4.53 | Q394 | 7.49 | G395 | 7.50 |
| C174 | 2.44 | R176 | 2.46 | W274 | 3.50 | P277 | 4.53 | Q394 | 7.49 | V398 | 7.53 |
| C174 | 2.44 | E408 | 8.49 | P277 | 4.53 | P312 | 5.42 | G395 | 7.50 | V398 | 7.53 |
| C174 | 2.44 | V409 | 8.50 | C296 | ECL2 | W297 | ECL2 | V398 | 7.53 | C403 | 7.58 |
| R176 | 2.46 | H180 | 2.50 | N320 | 5.50 | T353 | 6.42 | C403 | 7.58 | F404 | 7.59 |
| R176 | 2.46 | E247 | 3.50 | N320 | 5.50 | L356 | 6.45 | C403 | 7.58 | N406 | 8.47 |
| R176 | 2.46 | Y250 | 3.53 | N320 | 5.50 | P358 | 6.47 | C403 | 7.58 | E408 | 8.49 |
| R176 | 2.46 | L251 | 3.54 | N320 | 5.50 | L359 | 6.48 | C403 | 7.58 | V409 | 8.50 |
| R176 | 2.46 | E408 | 8.49 | N320 | 5.50 | G361 | 6.50 | F404 | 7.59 | N406 | 8.47 |
| H180 | 2.50 | W243 | 3.46 | T353 | 6.42 | L356 | 6.45 | F404 | 7.59 | V409 | 8.50 |
| H180 | 2.50 | E247 | 3.50 | T353 | 6.42 | P358 | 6.47 | N406 | 8.47 | E408 | 8.49 |
| H180 | 2.50 | V398 | 7.53 | T353 | 6.42 | G361 | 6.50 | N406 | 8.47 | V409 | 8.50 |
| H180 | 2.50 | C403 | 7.58 | L356 | 6.45 | P358 | 6.47 | E408 | 8.49 | V409 | 8.50 |
| C226 | 3.29 | C296 | ECL2 | | | | | | | | |

**Supplementary Table S8: Adaptive Sampling Metrics for GLP1R. Residue contacts between Residue 1 and Residue 2 were used as metrics for Adaptive Sampling. The Wootten numbering for each residue is reported as well.**

| Adaptive Sampling Metrics - PTH1R | | | | | | | | | | | |
|---|---|---|---|---|---|---|---|---|---|---|---|
| Residue 1 | W.N. 1 | Residue 2 | W.N. 2 | Residue 1 | W.N. 1 | Residue 2 | W.N. 2 | Residue 1 | W.N. 1 | Residue 2 | W.N. 2 |
| C108 | ECD | W118 | ECD | C281 | 3.29 | W352 | ECL2 | L413 | 6.45 | L416 | 6.48 |
| C108 | ECD | C148 | ECD | N295 | 3.43 | W298 | 3.46 | L413 | 6.45 | G418 | 6.50 |
| C108 | ECD | G152 | ECD | N295 | 3.43 | G328 | 4.49 | L413 | 6.45 | Q451 | 7.49 |
| D113 | ECD | C117 | ECD | N295 | 3.43 | W329 | 3.50 | L413 | 6.45 | V455 | 7.53 |
| D113 | ECD | W118 | ECD | N295 | 3.43 | L416 | 6.48 | P415 | 6.47 | L416 | 6.48 |
| C117 | ECD | W118 | ECD | N295 | 3.43 | G418 | 6.50 | P415 | 6.47 | G418 | 6.50 |
| C117 | ECD | C131 | ECD | W298 | 3.46 | E302 | 3.50 | P415 | 6.47 | F450 | 7.48 |
| C117 | ECD | P132 | ECD | W298 | 3.46 | G328 | 4.49 | P415 | 6.47 | Q451 | 7.49 |
| W118 | ECD | P132 | ECD | W298 | 3.46 | W329 | 3.50 | P415 | 6.47 | G452 | 7.50 |
| C131 | ECD | P132 | ECD | E302 | 3.50 | Y305 | 3.53 | P415 | 6.47 | V455 | 7.53 |
| C148 | ECD | G152 | ECD | E302 | 3.50 | L306 | 3.54 | L416 | 6.48 | G418 | 6.50 |
| G194 | 1.46 | S198 | 1.50 | E302 | 3.50 | N374 | 5.50 | L416 | 6.48 | F450 | 7.48 |
| G194 | 1.46 | F450 | 7.48 | E302 | 3.50 | T410 | 6.42 | L416 | 6.48 | Q451 | 7.49 |
| G194 | 1.46 | Q451 | 7.49 | E302 | 3.50 | L413 | 6.45 | L416 | 6.48 | G452 | 7.50 |
| G194 | 1.46 | G452 | 7.50 | E302 | 3.50 | L416 | 6.48 | L416 | 6.48 | V455 | 7.53 |
| S198 | 1.50 | L202 | 1.54 | E302 | 3.50 | G418 | 6.50 | G418 | 6.50 | F450 | 7.48 |
| S198 | 1.50 | F450 | 7.48 | Y305 | 3.53 | L306 | 3.54 | G418 | 6.50 | Q451 | 7.49 |
| S198 | 1.50 | Q451 | 7.49 | L306 | 3.54 | N374 | 5.50 | G418 | 6.50 | G452 | 7.50 |
| S198 | 1.50 | G452 | 7.50 | L306 | 3.54 | T410 | 6.42 | G418 | 6.50 | V455 | 7.53 |
| S198 | 1.50 | V455 | 7.53 | L306 | 3.54 | L413 | 6.45 | F450 | 7.48 | Q451 | 7.49 |
| L202 | 1.54 | G452 | 7.50 | L306 | 3.54 | L416 | 6.48 | F450 | 7.48 | G452 | 7.50 |
| L202 | 1.54 | V455 | 7.53 | G328 | 4.49 | W329 | 3.50 | F450 | 7.48 | V455 | 7.53 |
| L202 | 1.54 | C460 | 7.58 | G328 | 4.49 | P332 | 4.53 | Q451 | 7.49 | G452 | 7.50 |
| C217 | 2.44 | R219 | 2.46 | W329 | 3.50 | P332 | 4.53 | Q451 | 7.49 | V455 | 7.53 |
| C217 | 2.44 | E465 | 8.49 | P332 | 4.53 | P366 | 5.42 | G452 | 7.50 | V455 | 7.53 |
| C217 | 2.44 | V466 | 8.50 | C351 | ECL2 | W352 | ECL2 | V455 | 7.53 | C460 | 7.58 |
| R219 | 2.46 | H223 | 2.50 | N374 | 5.50 | T410 | 6.42 | C460 | 7.58 | F461 | 7.59 |
| R219 | 2.46 | E302 | 3.50 | N374 | 5.50 | L413 | 6.45 | C460 | 7.58 | N463 | 8.47 |
| R219 | 2.46 | Y305 | 3.53 | N374 | 5.50 | P415 | 6.47 | C460 | 7.58 | E465 | 8.49 |
| R219 | 2.46 | L306 | 3.54 | N374 | 5.50 | L416 | 6.48 | C460 | 7.58 | V466 | 8.50 |
| R219 | 2.46 | E465 | 8.49 | N374 | 5.50 | G418 | 6.50 | F461 | 7.59 | N463 | 8.47 |
| H223 | 2.50 | W298 | 3.46 | T410 | 6.42 | L413 | 6.45 | F461 | 7.59 | V466 | 8.50 |
| H223 | 2.50 | E302 | 3.50 | T410 | 6.42 | P415 | 6.47 | N463 | 8.47 | E465 | 8.49 |
| H223 | 2.50 | V455 | 7.53 | T410 | 6.42 | G418 | 6.50 | N463 | 8.47 | V466 | 8.50 |
| H223 | 2.50 | C460 | 7.58 | L413 | 6.45 | P415 | 6.47 | E465 | 8.49 | V466 | 8.50 |
| C281 | 3.29 | C351 | ECL2 | | | | | | | | |

**Supplementary Table S9: Adaptive Sampling Metrics for PTH1R. Residue contacts between Residue 1 and Residue 2 were used as metrics for Adaptive Sampling. The Wootten numbering for each residue is reported as well.**

| Adaptive Sampling Metrics - SCTR | | | | | | | | | | | |
|---|---|---|---|---|---|---|---|---|---|---|---|
| Residue 1 | W.N. 1 | Residue 2 | W.N. 2 | Residue 1 | W.N. 1 | Residue 2 | W.N. 2 | Residue 1 | W.N. 1 | Residue 2 | W.N. 2 |
| C66 | ECD | W76 | ECD | C215 | 3.29 | W286 | ECL2 | L347 | 6.45 | L350 | 6.48 |
| C66 | ECD | C107 | ECD | N229 | 3.43 | W232 | 3.46 | L347 | 6.45 | G352 | 6.50 |
| C66 | ECD | G111 | ECD | N229 | 3.43 | G262 | 4.49 | L347 | 6.45 | Q380 | 7.49 |
| D71 | ECD | C75 | ECD | N229 | 3.43 | W263 | 3.50 | L347 | 6.45 | V384 | 7.53 |
| D71 | ECD | W76 | ECD | N229 | 3.43 | L350 | 6.48 | P349 | 6.47 | L350 | 6.48 |
| C75 | ECD | W76 | ECD | N229 | 3.43 | G352 | 6.50 | P349 | 6.47 | G352 | 6.50 |
| C75 | ECD | C89 | ECD | W232 | 3.46 | E236 | 3.50 | P349 | 6.47 | F379 | 7.48 |
| C75 | ECD | P90 | ECD | W232 | 3.46 | G262 | 4.49 | P349 | 6.47 | Q380 | 7.49 |
| W76 | ECD | P90 | ECD | W232 | 3.46 | W263 | 3.50 | P349 | 6.47 | G381 | 7.50 |
| C89 | ECD | P90 | ECD | E236 | 3.50 | Y239 | 3.53 | P349 | 6.47 | V384 | 7.53 |
| C107 | ECD | G111 | ECD | E236 | 3.50 | L240 | 3.54 | L350 | 6.48 | G352 | 6.50 |
| G149 | 1.46 | S153 | 1.50 | E236 | 3.50 | N309 | 5.50 | L350 | 6.48 | F379 | 7.48 |
| G149 | 1.46 | F379 | 7.48 | E236 | 3.50 | T344 | 6.42 | L350 | 6.48 | Q380 | 7.49 |
| G149 | 1.46 | Q380 | 7.49 | E236 | 3.50 | L347 | 6.45 | L350 | 6.48 | G381 | 7.50 |
| G149 | 1.46 | G381 | 7.50 | E236 | 3.50 | L350 | 6.48 | L350 | 6.48 | V384 | 7.53 |
| S153 | 1.50 | L157 | 1.54 | E236 | 3.50 | G352 | 6.50 | G352 | 6.50 | F379 | 7.48 |
| S153 | 1.50 | F379 | 7.48 | Y239 | 3.53 | L240 | 3.54 | G352 | 6.50 | Q380 | 7.49 |
| S153 | 1.50 | Q380 | 7.49 | L240 | 3.54 | N309 | 5.50 | G352 | 6.50 | G381 | 7.50 |
| S153 | 1.50 | G381 | 7.50 | L240 | 3.54 | T344 | 6.42 | G352 | 6.50 | V384 | 7.53 |
| S153 | 1.50 | V384 | 7.53 | L240 | 3.54 | L347 | 6.45 | F379 | 7.48 | Q380 | 7.49 |
| L157 | 1.54 | G381 | 7.50 | L240 | 3.54 | L350 | 6.48 | F379 | 7.48 | G381 | 7.50 |
| L157 | 1.54 | V384 | 7.53 | G262 | 4.49 | W263 | 3.50 | F379 | 7.48 | V384 | 7.53 |
| L157 | 1.54 | C389 | 7.58 | G262 | 4.49 | P266 | 4.53 | Q380 | 7.49 | G381 | 7.50 |
| C172 | 2.44 | R174 | 2.46 | W263 | 3.50 | P266 | 4.53 | Q380 | 7.49 | V384 | 7.53 |
| C172 | 2.44 | E394 | 8.49 | P266 | 4.53 | P301 | 5.42 | G381 | 7.50 | V384 | 7.53 |
| C172 | 2.44 | V395 | 8.50 | C285 | ECL2 | W286 | ECL2 | V384 | 7.53 | C389 | 7.58 |
| R174 | 2.46 | H178 | 2.50 | N309 | 5.50 | T344 | 6.42 | C389 | 7.58 | F390 | 7.59 |
| R174 | 2.46 | E236 | 3.50 | N309 | 5.50 | L347 | 6.45 | C389 | 7.58 | N392 | 8.47 |
| R174 | 2.46 | Y239 | 3.53 | N309 | 5.50 | P349 | 6.47 | C389 | 7.58 | E394 | 8.49 |
| R174 | 2.46 | L240 | 3.54 | N309 | 5.50 | L350 | 6.48 | C389 | 7.58 | V395 | 8.50 |
| R174 | 2.46 | E394 | 8.49 | N309 | 5.50 | G352 | 6.50 | F390 | 7.59 | N392 | 8.47 |
| H178 | 2.50 | W232 | 3.46 | T344 | 6.42 | L347 | 6.45 | F390 | 7.59 | V395 | 8.50 |
| H178 | 2.50 | E236 | 3.50 | T344 | 6.42 | P349 | 6.47 | N392 | 8.47 | E394 | 8.49 |
| H178 | 2.50 | V384 | 7.53 | T344 | 6.42 | G352 | 6.50 | N392 | 8.47 | V395 | 8.50 |
| H178 | 2.50 | C389 | 7.58 | L347 | 6.45 | P349 | 6.47 | E394 | 8.49 | V395 | 8.50 |
| C215 | 3.29 | C285 | ECL2 | | | | | | | | |

**Supplementary Table S10: Adaptive Sampling Metrics for SCTR. Residue contacts between Residue 1 and Residue 2 were used as metrics for Adaptive Sampling. The Wootten numbering for each residue is reported as well.**

| Adaptive Sampling Metrics - PAC1R | | | | | | | | | | | |
|---|---|---|---|---|---|---|---|---|---|---|---|
| Residue 1 | W.N. 1 | Residue 2 | W.N. 2 | Residue 1 | W.N. 1 | Residue 2 | W.N. 2 | Residue 1 | W.N. 1 | Residue 2 | W.N. 2 |
| C54 | ECD | W64 | ECD | C226 | 3.29 | W297 | ECL2 | L358 | 6.45 | L361 | 6.48 |
| C54 | ECD | C118 | ECD | N240 | 3.43 | W243 | 3.46 | L358 | 6.45 | G363 | 6.50 |
| C54 | ECD | G122 | ECD | N240 | 3.43 | G273 | 4.49 | L358 | 6.45 | Q392 | 7.49 |
| D59 | ECD | C63 | ECD | N240 | 3.43 | W274 | 3.50 | L358 | 6.45 | V396 | 7.53 |
| D59 | ECD | W64 | ECD | N240 | 3.43 | L361 | 6.48 | P360 | 6.47 | L361 | 6.48 |
| C63 | ECD | W64 | ECD | N240 | 3.43 | G363 | 6.50 | P360 | 6.47 | G363 | 6.50 |
| C63 | ECD | C77 | ECD | W243 | 3.46 | E247 | 3.50 | P360 | 6.47 | F391 | 7.48 |
| C63 | ECD | P78 | ECD | W243 | 3.46 | G273 | 4.49 | P360 | 6.47 | Q392 | 7.49 |
| W64 | ECD | P78 | ECD | W243 | 3.46 | W274 | 3.50 | P360 | 6.47 | G393 | 7.50 |
| C77 | ECD | P78 | ECD | E247 | 3.50 | Y250 | 3.53 | P360 | 6.47 | V396 | 7.53 |
| C118 | ECD | G122 | ECD | E247 | 3.50 | L251 | 3.54 | L361 | 6.48 | G363 | 6.50 |
| G160 | 1.46 | S164 | 1.50 | E247 | 3.50 | N320 | 5.50 | L361 | 6.48 | F391 | 7.48 |
| G160 | 1.46 | F391 | 7.48 | E247 | 3.50 | T355 | 6.42 | L361 | 6.48 | Q392 | 7.49 |
| G160 | 1.46 | Q392 | 7.49 | E247 | 3.50 | L358 | 6.45 | L361 | 6.48 | G393 | 7.50 |
| G160 | 1.46 | G393 | 7.50 | E247 | 3.50 | L361 | 6.48 | L361 | 6.48 | V396 | 7.53 |
| S164 | 1.50 | L168 | 1.54 | E247 | 3.50 | G363 | 6.50 | G363 | 6.50 | F391 | 7.48 |
| S164 | 1.50 | F391 | 7.48 | Y250 | 3.53 | L251 | 3.54 | G363 | 6.50 | Q392 | 7.49 |
| S164 | 1.50 | Q392 | 7.49 | L251 | 3.54 | N320 | 5.50 | G363 | 6.50 | G393 | 7.50 |
| S164 | 1.50 | G393 | 7.50 | L251 | 3.54 | T355 | 6.42 | G363 | 6.50 | V396 | 7.53 |
| S164 | 1.50 | V396 | 7.53 | L251 | 3.54 | L358 | 6.45 | F391 | 7.48 | Q392 | 7.49 |
| L168 | 1.54 | G393 | 7.50 | L251 | 3.54 | L361 | 6.48 | F391 | 7.48 | G393 | 7.50 |
| L168 | 1.54 | V396 | 7.53 | G273 | 4.49 | W274 | 3.50 | F391 | 7.48 | V396 | 7.53 |
| L168 | 1.54 | C401 | 7.58 | G273 | 4.49 | P277 | 4.53 | Q392 | 7.49 | G393 | 7.50 |
| C183 | 2.44 | R185 | 2.46 | W274 | 3.50 | P277 | 4.53 | Q392 | 7.49 | V396 | 7.53 |
| C183 | 2.44 | E406 | 8.49 | P277 | 4.53 | P312 | 5.42 | G393 | 7.50 | V396 | 7.53 |
| C183 | 2.44 | V407 | 8.50 | C296 | ECL2 | W297 | ECL2 | V396 | 7.53 | C401 | 7.58 |
| R185 | 2.46 | H189 | 2.50 | N320 | 5.50 | T355 | 6.42 | C401 | 7.58 | F402 | 7.59 |
| R185 | 2.46 | E247 | 3.50 | N320 | 5.50 | L358 | 6.45 | C401 | 7.58 | N404 | 8.47 |
| R185 | 2.46 | Y250 | 3.53 | N320 | 5.50 | P360 | 6.47 | C401 | 7.58 | E406 | 8.49 |
| R185 | 2.46 | L251 | 3.54 | N320 | 5.50 | L361 | 6.48 | C401 | 7.58 | V407 | 8.50 |
| R185 | 2.46 | E406 | 8.49 | N320 | 5.50 | G363 | 6.50 | F402 | 7.59 | N404 | 8.47 |
| H189 | 2.50 | W243 | 3.46 | T355 | 6.42 | L358 | 6.45 | F402 | 7.59 | V407 | 8.50 |
| H189 | 2.50 | E247 | 3.50 | T355 | 6.42 | P360 | 6.47 | N404 | 8.47 | E406 | 8.49 |
| H189 | 2.50 | V396 | 7.53 | T355 | 6.42 | G363 | 6.50 | N404 | 8.47 | V407 | 8.50 |
| H189 | 2.50 | C401 | 7.58 | L358 | 6.45 | P360 | 6.47 | E406 | 8.49 | V407 | 8.50 |
| C226 | 3.29 | C296 | ECL2 | | | | | | | | |

**Supplementary Table S11: Adaptive Sampling Metrics for PAC1R. Residue contacts between Residue 1 and Residue 2 were used as metrics for Adaptive Sampling. The Wootten numbering for each residue is reported as well.**

| Adaptive Sampling Metrics - CALCR | | | | | | | | | | | |
|---|---|---|---|---|---|---|---|---|---|---|---|
| Residue 1 | W.N. 1 | Residue 2 | W.N. 2 | Residue 1 | W.N. 1 | Residue 2 | W.N. 2 | Residue 1 | W.N. 1 | Residue 2 | W.N. 2 |
| C72 | ECD | W82 | ECD | C219 | 3.29 | W290 | ECL2 | L348 | 6.45 | L351 | 6.48 |
| C72 | ECD | C112 | ECD | N233 | 3.43 | W236 | 3.46 | L348 | 6.45 | G353 | 6.50 |
| C72 | ECD | G116 | ECD | N233 | 3.43 | G266 | 4.49 | L348 | 6.45 | Q383 | 7.49 |
| D77 | ECD | C81 | ECD | N233 | 3.43 | W267 | 3.50 | L348 | 6.45 | V387 | 7.53 |
| D77 | ECD | W82 | ECD | N233 | 3.43 | L351 | 6.48 | P350 | 6.47 | L351 | 6.48 |
| C81 | ECD | W82 | ECD | N233 | 3.43 | G353 | 6.50 | P350 | 6.47 | G353 | 6.50 |
| C81 | ECD | C95 | ECD | W236 | 3.46 | E240 | 3.50 | P350 | 6.47 | F382 | 7.48 |
| C81 | ECD | P96 | ECD | W236 | 3.46 | G266 | 4.49 | P350 | 6.47 | Q383 | 7.49 |
| W82 | ECD | P96 | ECD | W236 | 3.46 | W267 | 3.50 | P350 | 6.47 | G384 | 7.50 |
| C95 | ECD | P96 | ECD | E240 | 3.50 | Y243 | 3.53 | P350 | 6.47 | V387 | 7.53 |
| C112 | ECD | G116 | ECD | E240 | 3.50 | L244 | 3.54 | L351 | 6.48 | G353 | 6.50 |
| G155 | 1.46 | S159 | 1.50 | E240 | 3.50 | N312 | 5.50 | L351 | 6.48 | F382 | 7.48 |
| G155 | 1.46 | F382 | 7.48 | E240 | 3.50 | T345 | 6.42 | L351 | 6.48 | Q383 | 7.49 |
| G155 | 1.46 | Q383 | 7.49 | E240 | 3.50 | L348 | 6.45 | L351 | 6.48 | G384 | 7.50 |
| G155 | 1.46 | G384 | 7.50 | E240 | 3.50 | L351 | 6.48 | L351 | 6.48 | V387 | 7.53 |
| S159 | 1.50 | L163 | 1.54 | E240 | 3.50 | G353 | 6.50 | G353 | 6.50 | F382 | 7.48 |
| S159 | 1.50 | F382 | 7.48 | Y243 | 3.53 | L244 | 3.54 | G353 | 6.50 | Q383 | 7.49 |
| S159 | 1.50 | Q383 | 7.49 | L244 | 3.54 | N312 | 5.50 | G353 | 6.50 | G384 | 7.50 |
| S159 | 1.50 | G384 | 7.50 | L244 | 3.54 | T345 | 6.42 | G353 | 6.50 | V387 | 7.53 |
| S159 | 1.50 | V387 | 7.53 | L244 | 3.54 | L348 | 6.45 | F382 | 7.48 | Q383 | 7.49 |
| L163 | 1.54 | G384 | 7.50 | L244 | 3.54 | L351 | 6.48 | F382 | 7.48 | G384 | 7.50 |
| L163 | 1.54 | V387 | 7.53 | G266 | 4.49 | W267 | 3.50 | F382 | 7.48 | V387 | 7.53 |
| L163 | 1.54 | C392 | 7.58 | G266 | 4.49 | P270 | 4.53 | Q383 | 7.49 | G384 | 7.50 |
| C178 | 2.44 | R180 | 2.46 | W267 | 3.50 | P270 | 4.53 | Q383 | 7.49 | V387 | 7.53 |
| C178 | 2.44 | E397 | 8.49 | P270 | 4.53 | P304 | 5.42 | G384 | 7.50 | V387 | 7.53 |
| C178 | 2.44 | V398 | 8.50 | C289 | ECL2 | W290 | ECL2 | V387 | 7.53 | C392 | 7.58 |
| R180 | 2.46 | H184 | 2.50 | N312 | 5.50 | T345 | 6.42 | C392 | 7.58 | F393 | 7.59 |
| R180 | 2.46 | E240 | 3.50 | N312 | 5.50 | L348 | 6.45 | C392 | 7.58 | N395 | 8.47 |
| R180 | 2.46 | Y243 | 3.53 | N312 | 5.50 | P350 | 6.47 | C392 | 7.58 | E397 | 8.49 |
| R180 | 2.46 | L244 | 3.54 | N312 | 5.50 | L351 | 6.48 | C392 | 7.58 | V398 | 8.50 |
| R180 | 2.46 | E397 | 8.49 | N312 | 5.50 | G353 | 6.50 | F393 | 7.59 | N395 | 8.47 |
| H184 | 2.50 | W236 | 3.46 | T345 | 6.42 | L348 | 6.45 | F393 | 7.59 | V398 | 8.50 |
| H184 | 2.50 | E240 | 3.50 | T345 | 6.42 | P350 | 6.47 | N395 | 8.47 | E397 | 8.49 |
| H184 | 2.50 | V387 | 7.53 | T345 | 6.42 | G353 | 6.50 | N395 | 8.47 | V398 | 8.50 |
| H184 | 2.50 | C392 | 7.58 | L348 | 6.45 | P350 | 6.47 | E397 | 8.49 | V398 | 8.50 |
| C219 | 3.29 | C289 | ECL2 | | | | | | | | |

**Supplementary Table S12: Adaptive Sampling Metrics for CALCR. Residue contacts between Residue 1 and Residue 2 were used as metrics for Adaptive Sampling. The Wootten numbering for each residue is reported as well.**

(6) Zhao, L.-H.; Ma, S.; Sutkeviciute, I.; Shen, D.-D.; Zhou, X. E.; de Waal, P. W.; Li, C.-

Y.; Kang, Y.; Clark, L. J.; Jean-Alphonse, F. G.; White, A. D.; Yang, D.; Dai, A.; Cai, X.; Chen, J.; Li, C.; Jiang, Y.; Watanabe, T.; Gardella, T. J.; Melcher, K.; Wang, M.-W.; Vilardaga, J.-P.; Xu, H. E.; Zhang, Y. Structure and dynamics of the active human parathyroid hormone receptor-1. *Science* **2019**, *364*, 148–153.
